# Rubber mapping reveals opportunities and current limitations of deforestation due-diligence

**DOI:** 10.64898/2026.07.25.740673

**Authors:** A. Ahrends, S. B. Harrison, P. M. Hollingsworth, J. D. J. Heath, Y. Wang, J. Xu, J. M. H. Green

**Author notes:** These authors contributed equally to the production of the work (A. Ahrends, S. B. Harrison).

## Abstract

Commodity maps and the ability to monitor commodity-driven deforestation are essential for sustainability risk assessment and due diligence. Natural rubber (*Hevea brasiliensis*), used primarily in tyres, remains challenging to map because of its similarity to other tree cover. Integrating optical and radar satellite data, we produced a 10 m map of rubber distribution in Southeast Asia for around 2020 and assessed this map and similar products for deforestation due diligence. In continental Southeast Asia, estimated user’s accuracy was 0.95 and area-adjusted producer’s accuracy 0.78. Rubber-associated clearances of natural and semi-natural tree cover during 2011-2016 were estimated at 0.3-0.4 Mha, predominantly associated with industrial-scale plantations and concentrated in Cambodia. Historical clearances were less certain, with plausible bounds of 1.3-3.0 Mha since 1990. Evaluation of rubber and forest maps highlighted persistent limitations, particularly in insular Southeast Asia and smallholder systems. Accounting for these limitations is critical for fair and effective due diligence.

## Introduction

Natural rubber, from the tree *Hevea brasiliensis* (Willd. ex A.Juss.) Müll.Arg., is widely used in tyres and other products. Its properties are difficult to replicate^1^ and synthetic alternatives are energy- and carbon-intensive^2^. In contrast, natural rubber can sequester carbon^3^ and provide income for smallholders, who account for over 80% of global production^4^, which now approaches 15 Mt annually^5^.

Rubber cultivation is, however, also associated with social^6,7^ and environmental concerns^8,9^, including deforestation^10–14^. In response, many companies have adopted sustainability commitments^15,16^, and regulations such as the EU Deforestation Regulation (EUDR)^17^ aim to mandate due diligence in supply chains. These efforts require spatial production data and maps that distinguish rubber from natural forest. This is challenging because of rubber’s spectral similarity to other tree cover^18^. Existing datasets either cover limited areas^19–24^ or require refinement to minimise confusion between rubber and other tree cover, as in the first region-wide map by Wang et al^25–27^. A recent pantropical probability-based rubber map^28^, derived from geospatial foundation-model embeddings, substantially extends spatial coverage. However, based on opportunistic training and validation data, its performance has not yet been systematically assessed^28^. At the same time, global forest maps^29–34^ can misclassify rubber as natural forest, complicating deforestation monitoring^35^.

Partly owing to the difficulty of mapping rubber, published estimates of rubber-linked deforestation vary by over an order of magnitude. The highest estimate suggests over 5 Mha since 2003 in continental Southeast Asia alone^36^. Wang et al.^25^ estimated 2.5 Mha across Southeast Asia since 2000, but noted that, due to challenges in distinguishing planted from natural tree cover, this figure can include the conversion or rotation of older plantations and agroforests, including rubber itself. Sheil et al.^27^ argued that the “actual extent” of deforestation was only one-quarter (0.6 Mha); however, their analysis was restricted to primary humid forest, thereby excluding deciduous and secondary forests^18^.

Given the need for baseline rubber maps with well-understood accuracy and minimal conflation with forest, and a more nuanced understanding of rubber impacts and risk geographies, here we:

1. Develop a new 10 m map of mature rubber plantations for 2020 across Southeast Asia and explore the utility of this and other recent rubber and forest maps for deforestation due-diligence, focusing on conflation between rubber and natural forest.
2. Assess where mature rubber mapped for 2020 was associated with clearances during 2011-2016, the latest window for clearances contributing to mature rubber in 2020.
3. Approximate the scale of historical rubber-associated clearances of natural and seminatural tree cover from 1990 to 2016.

For objectives 2 and 3, clearance estimates were derived primarily through visual interpretation of satellite imagery at sample points to distinguish plantation rotation from losses of non-plantation tree cover. We refer to the latter as “natural and semi-natural tree cover”, including primary, secondary and degraded forests in both humid and seasonally dry systems. Semi-natural tree cover denotes naturally regenerating tree cover with visible human influence, such as prior clearing, grazing, and/or logging. Clearances described as “associated with” rubber refer to post-clearance land cover and do not imply causation.

## Results

### New high-resolution map of rubber

We mapped 8.54 Mha of rubber (Fig. 1), with an error-adjusted estimate of 12.53 ± 1.65 Mha (Table S1), suggesting that the mapped area is conservative. Estimated areas broadly align with official and subnational data, except in China, Myanmar and Laos, where our estimates are higher (Tables 1 and S2; Fig. S1).

**Figure 1.**
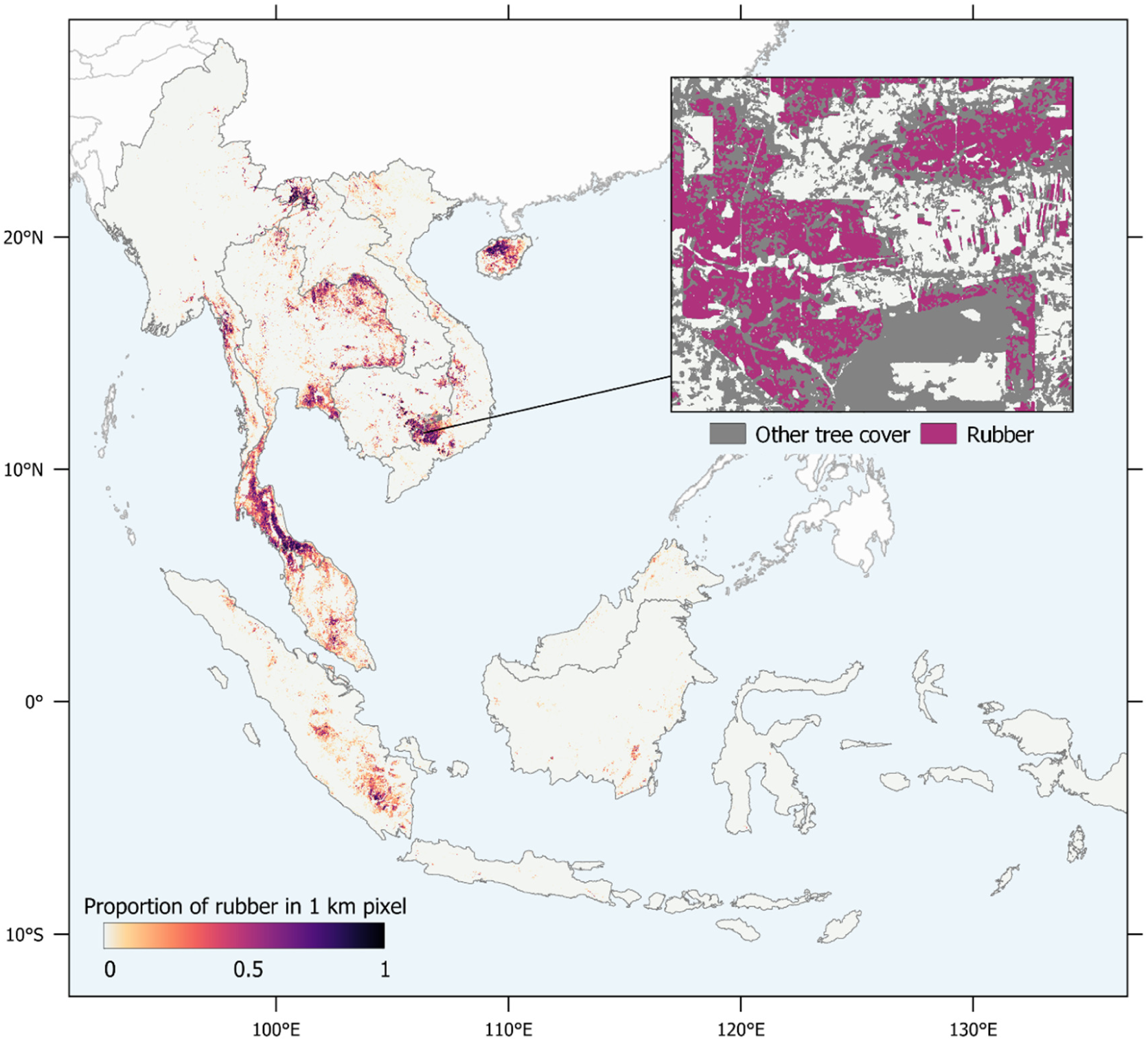
| Mapped rubber in ∼2020. Mapped rubber is aggregated to 1-km grid cells as the proportion of underlying 10 × 10 m rubber pixels. The inset shows the 10 m classification, which distinguishes rubber from other tree cover. Accuracy was higher in continental than insular Southeast Asia, especially for false negatives.

Validation against 2,500 sample points yielded an area-adjusted^37^ overall accuracy of 0.98, with relatively low commission (false rubber-positive) errors (user’s accuracy, UA=0.90) but more substantial omission (false rubber-negative) errors (producer’s accuracy, PA=0.63). Omission errors were concentrated in Malaysia and Indonesia (hereafter “insular Southeast Asia”), with higher rubber detection across the rest of the study area (“continental Southeast Asia”; UA=0.95; PA=0.78; Tables S3-S4). Sensitivity analyses treating cases of interpreter uncertainty as map errors primarily increased omission errors, suggesting that total rubber area is unlikely to be overestimated but may be underestimated (Tables S1, S3-S4). Residual commission errors included confusion with other plantations, open forest types, clearings, and young plantations, where bare ground in the dry season followed by green-up during the rainy season can resemble rubber phenology (Table S5). Commissions also resulted from a focal majority filter (see Methods), classifying small non-rubber patches embedded within rubber stands as rubber.

### Strengths and limitations of current maps for deforestation due diligence

We assessed strengths and limitations of widely-used rubber and forest maps for due diligence in rubber supply chains by quantifying classification errors that can obscure forest loss or generate false-positive deforestation alerts.

For forest we assessed forest omission and rubber conflation across 19 maps. Except for maps restricted to humid forest, omission rates were often below 10%. They mainly affected deciduous and semi-evergreen forests. Earlier datasets had substantial conflation with rubber; e.g. in the case of the first EC JRC Global Forest Cover 2020 map^38^ over 50% of the area mapped as rubber here was classified as natural forest (Table S6). In recent datasets, separation from rubber has improved, but false positives remain, particularly in insular Southeast Asia.

A Thailand national forest map^39^, the only national dataset available to us, performed best. Among semi-global forest datasets, recent Joint Research Centre^31,33^ and Google DeepMind/Nature Trace^30,34^ products had the best trade-off between minimising forest omission and rubber conflation (results for all datasets are in Table S7).

Analysis of 12 rubber products^19,22–25,28,36,40–42^ highlighted low detection power in Malaysia and Indonesia for all, except a local Kalimantan map^19^ (Table S8). This map also covers rubber agroforest; by contrast, our and other region-wide and pantropical rubber maps almost exclusively capture monocultures. Agroforests with mixed-species canopies were largely undetected and, in global forest datasets, often classified as natural forest (Table S9).

Where rubber maps conflated forest, this mainly affected deciduous forests. Compared with other region-wide products, our map had lower commission error (Table S9). However, because this comparison partly used data that trained our model, it should be interpreted with caution.

### Estimated scale of clearances in early 2010s

To estimate rubber-associated clearances of (semi-)natural tree cover between 2011 and 2016, we quantified the overlap between areas mapped as rubber in ∼2020 here and areas mapped as natural forest in ∼2011 in forest maps. We found only two historical forest maps that exclude rubber from “forest”: for primary humid evergreen forest^43^ and dry deciduous dipterocarp^44^ forests. As these maps only cover subsets of regional forest types, we also applied the LandTrendr algorithm^45^ to detect losses elsewhere.

To account for false clearance positives, we visually verified random samples from mapped overlap between rubber and forest layers using very-high-resolution imagery. Corresponding downward adjustments (Table S10) yielded an estimated clearance of 0.3 Mha. To account for rubber omission and commission errors in our map, we also calibrated this area by national rubber areas, yielding an estimate of ∼0.4 Mha. However, much of the increase was contributed by Indonesia where uncertainty was highest. We thus estimate clearances to lie within the range of 0.3-0.4 Mha (Table 2).

**Table 1.** | Rubber area in Southeast Asia according to this and previous studies. Discrepancies in country-level estimates across studies illustrate the challenges of mapping rubber. In the case of the probability-based Forest Data Partnership products areas also depend on threshold choice. Here we applied thresholds reported to balance precision and recall in global FDP evaluations. This results in over-mapping in Laos, Myanmar, and Vietnam, where according to our evaluations there is a risk of conflation with deciduous forests, and under-mapping in Indonesia, where detection power is low. As noted by the FDP authors^28^, there is a need for local accuracy assessments to select the best thresholds. A comparison of additional rubber maps is provided in Table S13. For China, only the main rubber-producing areas are included: Xishuangbanna and Hainan.

| (kha) | Cambodia | China | Indonesia | Laos | Malaysia | Myanmar | Thailand | Vietnam | Total |
| --- | --- | --- | --- | --- | --- | --- | --- | --- | --- |
| This study: error-adjusted estimate (mapped area) | 378<br>(392) | 1,256<br>(952) | 3,705<br>(1,200) | 385<br>(228) | 949<br>(867) | 907<br>(621) | 4,040<br>(3,436) | 878<br>(849) | **12,497**<br>(8,544) |
| Official rubber area<sup>‡</sup> | 404 | 813 | 3,776 | 275 | 1,137 | 659 | 3,915 | 931 | **11,915** |
| Wang et al.<sup>25</sup>: mapped area | 615 | 1,069 | 4,582 | 589 | 989 | 789 | 3,706 | 1,612 | **13,951** |
| Forest Data Partnership<sup>28</sup> model b: mapped area<sup>§</sup> | 751<br>985 | 1,392<br>1,583 | 90<br>113 | 2,163<br>3,049 | 491<br>563 | 3,315<br>4,675 | 5,687<br>6,615 | 3,303<br>4,330 | **17,192**<br>**21,914** |
| Forest Data Partnership<sup>22</sup> model a: mapped area | - | 957<br>1,258 | 2,692<br>4,837 | - | 396<br>703 | - | 3,048<br>5,099 | 2,281<br>3,732 | - |
| De Keersmaecker et al.<sup>42</sup>: mapped area | 413 | 534 | 1,301 | 868 | 263 | 1,148 | 2,952 | 657 | **8,135** |
| SDPT<sup>40,47</sup> version 3 (version 2): mapped area<sup>**</sup> | 858<br>(866) | - | 1,115<sup>††</sup><br>(1,115) | 686<br>(706) | 421<br>(421) | 603<br>(603) | 4,223<br>(4,250) | 587<br>(591) | - |

<sup>‡</sup> The official areas are not always directly comparable. Areas for Thailand and Indonesia refer to mature rubber in 2021, for Vietnam to harvested rubber in 2021, for Cambodia to planted rubber in 2021, and for Laos and Myanmar to planted rubber in 2018/19. Areas for China are for 2022. Sources of official data are reported in Table S13.
<sup>§</sup> The Forest Data Partnership (FDP) rubber probability models have been thresholded at two values reported to balance precision and recall in FDP evaluations: 0.59/0.44 and 0.23/0.35 for models 2025a/2025b, respectively.
<sup>\*\*</sup> Sources contributing to SDPT include Petersen et al. (2016), Xiao et al. (2021), Condro et al. (2020) and Debonne et al. (2019). Petersen et al. (2016) also mapped rubber mixed with other crops in Malaysia, Indonesia and Cambodia, and Debonne et al. (2019) did so for Cambodia. For comparability, we included only pure rubber areas. SDPT version 3 is under review and changes are still possible. In version 3, non-rubber datasets have been added for Cambodia, Laos, Myanmar, Thailand and Vietnam. These take precedence over Xiao et al. (2021) and Debonne et al. (2019), reducing rubber extent in areas of overlap (J. Richter, personal communication, 2026).
<sup>††</sup> Figure is approximate.

**Table 2.** | Estimated loss of natural and semi-natural tree cover associated with rubber between 2011 and 2016. Estimated losses of primary forest (PF), dry dipterocarp forest (DF), and other tree cover were adjusted for false positives using validation of 600 sample points (Table S10). The summed adjusted losses were then extrapolated by scaling the proportion of mapped rubber area estimated to have experienced losses to the total rubber area estimated in this study (a) and reported in official statistics (b). Estimated losses and especially extrapolated values should be interpreted with caution. Losses may be overestimated where large parts of the rubber area were estimated rather than directly mapped (e.g. in Indonesia), and underestimated where large clearances were underrepresented by random sample points (potentially Cambodia). The dry dipterocarp forest layer was not available for all countries.

| (ha) | PF<br>adjusted | DF<br>adjusted | Other<br>adjusted | Total<br>adjusted | %<br>mapped | Extrapolated<br>loss (a) | Extrapolated<br>loss (b) |
| --- | --- | --- | --- | --- | --- | --- | --- |
| Cambodia | 67,704 | 5,800 | 44,768 | **118,273** | 30 | *114,049* | *122,035* |
| China | 912 | NA | 0 | **912** | 0 | *1,203* | *779* |
| Indonesia | 18,866 | NA | 16,072 | **34,938** | 3 | *107,840* | *109,928* |
| Laos | 12,092 | 976 | 7,466 | **20,534** | 9 | *34,659* | *24,775* |
| Malaysia | 29,388 | NA | 4,641 | **34,029** | 4 | *37,252* | *44,650* |
| Myanmar | 4,709 | 12,063 | 29,937 | **46,710** | 8 | *68,238* | *49,614* |
| Thailand | 3,397 | 1,038 | 11,449 | **15,884** | 0 | *18,679* | *18,097* |
| Vietnam | 22,721 | 3,984 | 11,492 | **38,197** | 4 | *39,527* | *41,869* |
|  | **159,790** | **23,861** | **125,825** | **309,476** | **4** | ***421,446*** | ***411,747*** |

With respect to risk geographies, we detected extensive clearances in Cambodia, where large concession areas have been awarded for rubber^46^ (Fig. 2). Larger clearances also occurred in Myanmar, Vietnam, Malaysia, and Laos. Evidence for post-2010 clearances in Thailand and China was limited; estimates for Indonesia require caution (above). While clearances in Cambodia and Vietnam often affected natural tree cover, there was also widespread conversion of semi-natural tree cover, for example, in Myanmar and Thailand (Fig. 3, Table 2).

**Figure 2.**
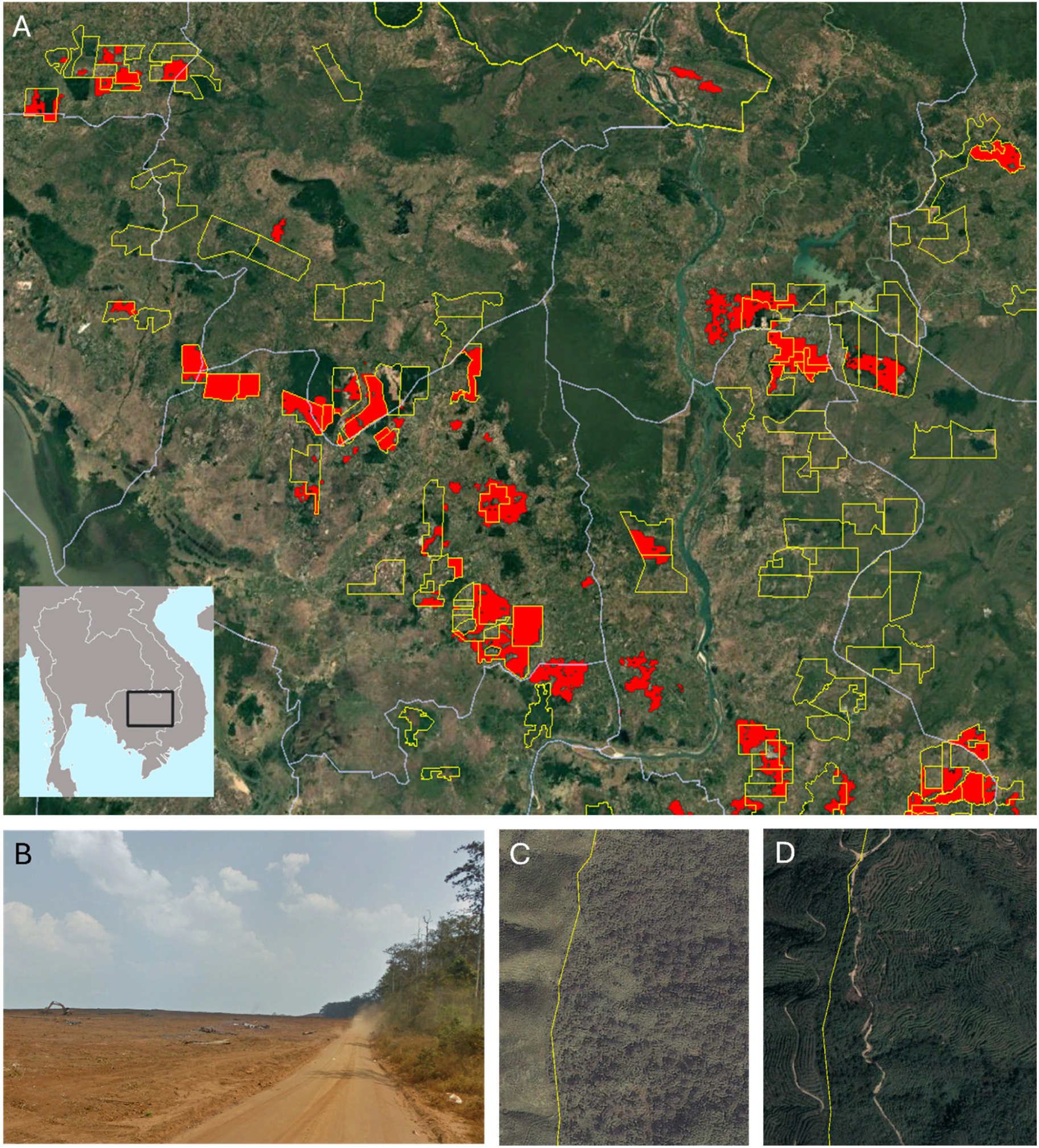
| Examples of clearances potentially associated with rubber. A: There was evidence of extensive clearance for industrial rubber production in Cambodia (red polygons), both inside and outside (mixed) rubber concession areas^46^ (yellow polygons). Several concessions still had forest in 2025, including extensive areas of deciduous forests (e.g. lower right in the image). Areas shown as deforested have been hand-digitised using very-high resolution imagery and may include interpretation errors. *Imagery: Landsat/Copernicus*. **B:** Example of clearance preceding rubber cultivation (*Google Street View imagery* at 107.529°E, 12.435°N from March 2014). **C:** Seminatural tree cover in Myanmar with smallholder fields in 2001 (east of the China-Myanmar border in yellow). D: The same area subsequently converted to industrial-scale rubber by 2018. *Imagery: Maxar Technologies*.

**Figure 3.**
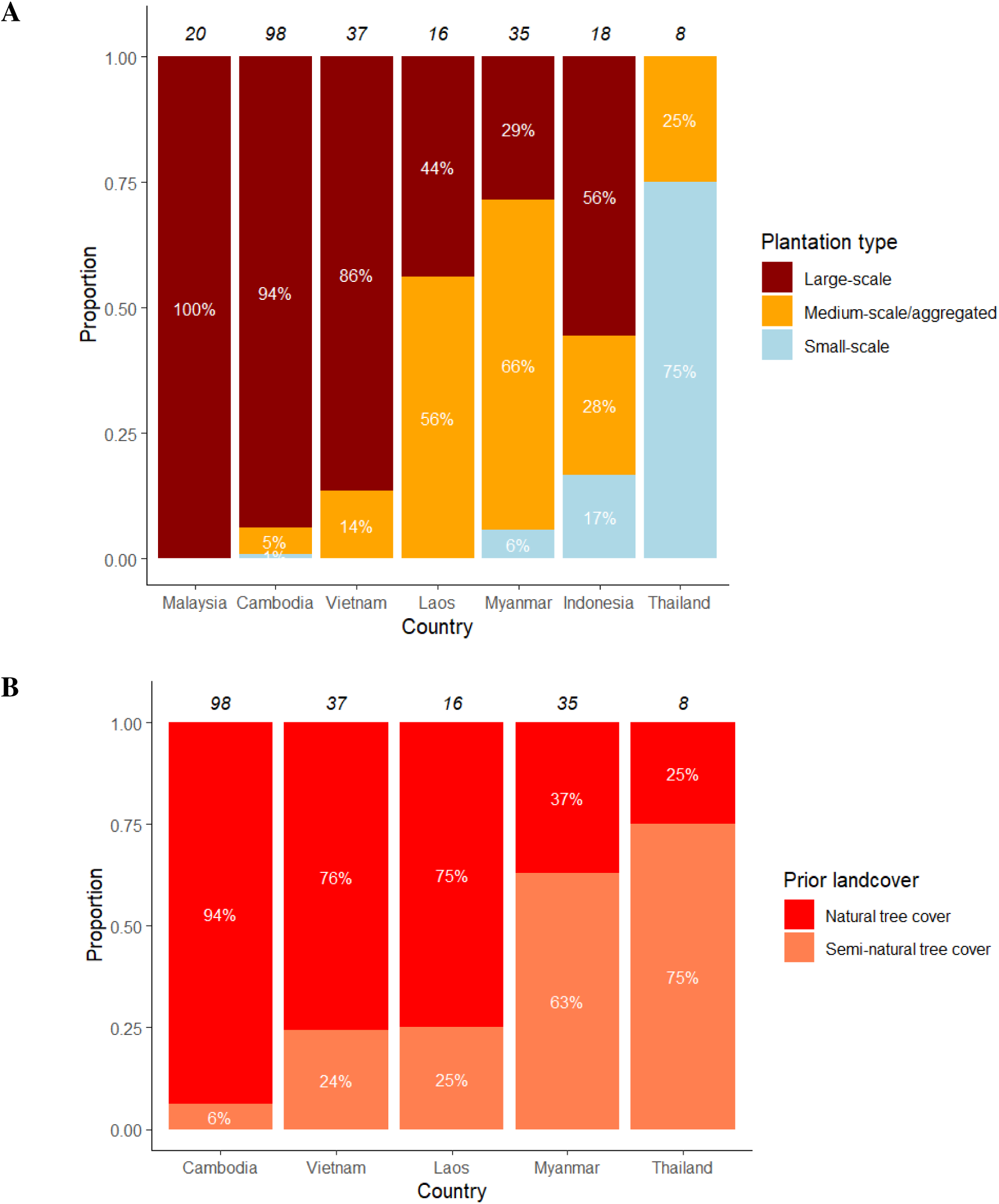
| Scale of rubber plantations (A) and types of tree cover (B) associated with clearances 2011-2016. Large-scale refers to estates typically >500 ha in size and characterised by uniform and often grid-style planting; medium-scale/aggregated to contiguous rubber landscapes composed of numerous aggregated smaller plantations, typical of e.g. farmer-investor arrangements; and small-scale to plantations <5 ha in size embedded within heterogenous land cover or natural vegetation. Separating semi-natural and natural tree cover (B) requires highquality imagery and was therefore not possible for Malaysia and Indonesia. For China the number of interpretable points was too low.

Industrial-scale plantations (Fig. 3) accounted for 74% of the estimated clearance area. A further 21% was associated with large clusters of smaller plantations, typical e.g. of investor-farmer arrangements. Only 5% was associated with small-scale rubber production.

### Approximate scale of historical clearances

To approximate the scale of historical clearances potentially associated with rubber, we visually interpreted land cover trajectories for 586 random rubber samples. The results are best interpreted as a bounded rather than point estimate as historical image quality, interpreter uncertainty, and challenging-to-interpret land-cover often limited precise reconstructions. Classification uncertainty was highest for earlier time periods and in equatorial regions (Fig. S3).

Around two-thirds of the point samples were located on land already modified by 1990. For 17% land-cover trajectories could not be determined. The remaining 17% were potentially associated with clearances; 60% affecting natural and 40% semi-natural tree cover. Around 10-15% of clearances affected areas with signs of heavy degradation, such as fragmented forest patches in poor condition or previously logged areas now dominated by scrub.

Extrapolation based on these proportions suggested ∼2 Mha of clearance since 1990 (1.7-2.4 Mha using the 95% bounds of estimated rubber area), and a similar area with unknown land-cover trajectory. The unknown area was larger when estimated by country rather than across the region, reflecting substantial uncertainty, particularly for Indonesia (Fig. 3; Table S11). To account for unknown trajectories and interpreter variability, we repeated the extrapolation using the most and least conservative interpretations across interpreters. This yielded a broad range of 1.3-3.0 Mha (Table S11), suggesting that clearance associated with rubber since 1990 was likely on the order of one to three million hectares, subject to forest definitions and interpretation uncertainty.

In long-established rubber-production areas (Thailand, Indonesia and Malaysia), many current rubber areas were already rubber or otherwise modified by 1990. In contrast, in Cambodia and Laos there was frequent evidence of clearances. Trajectories in China, Vietnam and Myanmar were heterogeneous. Where there were clearances, these often occurred earlier in the 1990s or 2000s; however, in Vietnam and Myanmar, several larger clearances also occurred relatively recently in the early-to-mid 2010s (Fig. 4; Fig. S2). Proportions of (mapped) rubber within protected areas and/or Key Biodiversity Areas were highest in Cambodia, Vietnam and Laos, ranging from 7% to 10%. The regional average was 4% (Table S12).

**Figure 4.**
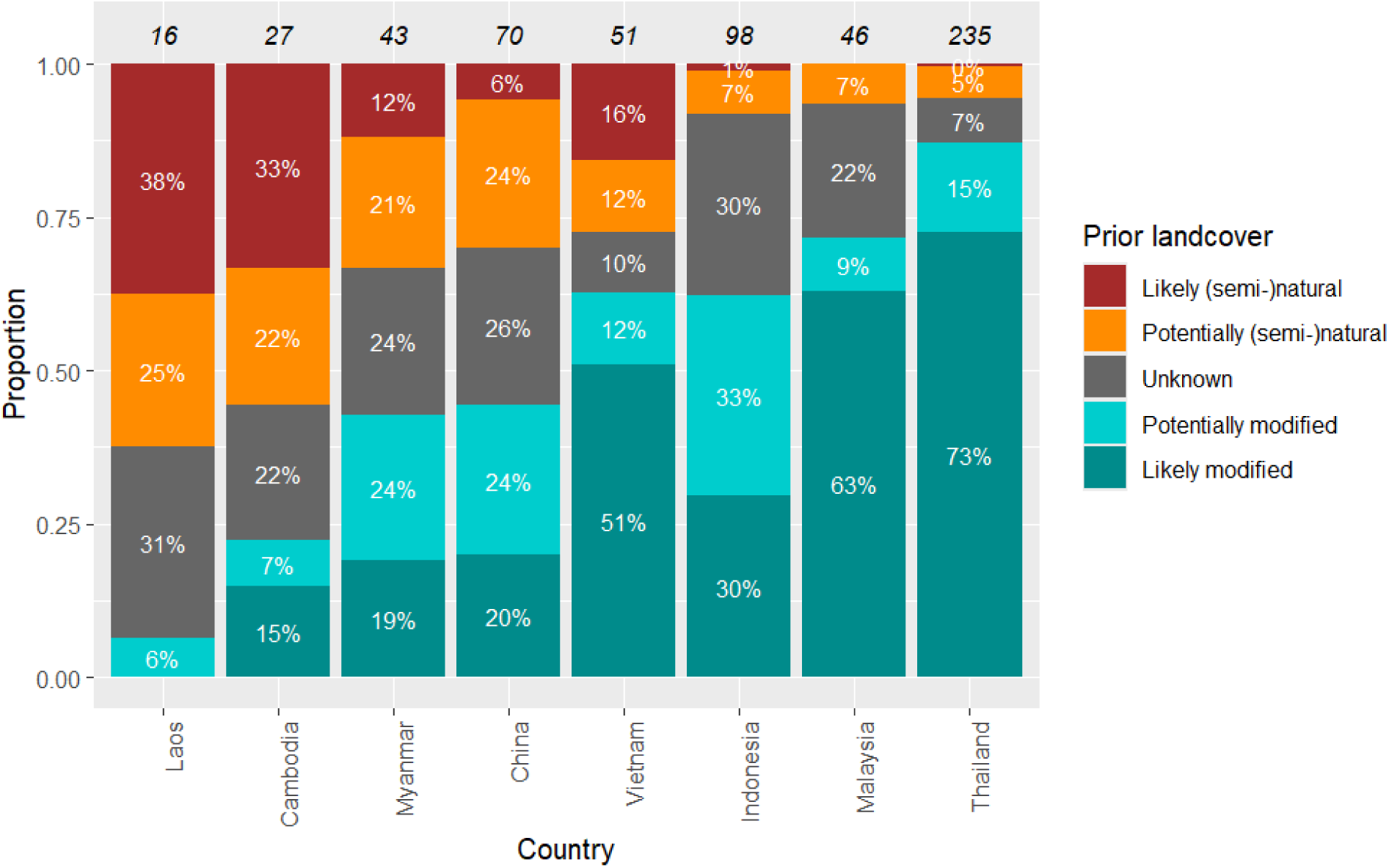
| Land cover prior to rubber cultivation for 586 reference points. The baseline for this analysis is 1990. Interpreters subjectively assigned certainty scores, ranging from 0.75 (highly uncertain) to 1 (virtually certain). Points that are ‘likely’ natural or modified received a certainty score of 0.85 or above, and points that are ‘potentially’ natural or modified had lower certainty. Points that interpreters felt unable to class were assigned to ‘unknown’.

## Discussion

Here, we present a conservative 10 m map of mature rubber plantations across Southeast Asia and combine it with analysis of land-cover trajectories to estimate rubber-associated clearances. We also assess what current rubber and forest maps can contribute to due diligence, and where limitations lie. Evaluation of our map suggested comparatively low forest conflation, an important requirement for due diligence. Our estimated rubber areas generally aligned with official statistics, except in China, Laos and Myanmar, where they are higher and may require caution. The new map, algorithm and input data are publicly available.

### Potential and limitations for use in policy

In practice, the main area of use we see for the map is to support supply chain due diligence and risk screening, for example in the context of EUDR and particularly in continental Southeast Asia where accuracy is higher. Pilot use with operators suggests that the map can provide additional evidence for supplier-mapped rubber areas by confirming that tree cover exhibits spectral characteristics consistent with rubber. However, due to the constraints outlined below the converse is not true - an absence of rubber in our map, particularly in regions (Malaysia and Indonesia) and systems (non-monoculture) where detection is poor, does not imply absence of rubber.

Several limitations should be considered. First, our accuracy statistics relied on image interpretation rather than field validation. Second, although our evaluation indicates comparatively low levels of conflation, some confusion between rubber and other tree cover, including forest, remains. Third, both in our and other region-wide rubber maps^22,25,28,36,40,42,47^, omission errors remain substantial in parts of insular Southeast Asia due to cloud cover, highly heterogenous land cover and weaker phenological signals. Finally, our algorithm partly relies on rubber’s characteristically rapid seasonal leaf-loss and renewal cycle, which emerges once trees reach approximately five years. Young plantations and diseased stands are therefore not detected. Detection may also be lower for plantations where undergrowth has not been cleared.

An added challenge - particularly relevant for smallholder rubber in Indonesia - are plantations with mixed-species canopies and multiple strata (“jungle” or agroforestry rubber). Unless rubber dominates the canopy, these systems are largely undetected in our and other region-wide rubber maps. This is not necessarily a failure: treating these systems separately from monoculture rubber is useful as their conversion to monocultures can constitute a loss of structural and biological diversity. However, because agroforests are frequently not distinguished from forest in natural forest maps, dedicated mapping efforts such as by le Maire et al.^19^ are required.

### Practical recommendations for due diligence

We recommend using several sources of evidence. Where available, local field data are best. Provided limitations are recognised, satellite-based products such as our and other publicly accessible maps, for example by the Forest Data Partnership^22,28^, De Keersmaecker et al.^42^ and le Maire et al.^19^, can add evidence. These products have different strengths and limitations. For example, our map was systematically validated, but only covers Southeast Asia and prioritises conservative classification over full detection. Recent pantropical maps^28^ have broader coverage, and, in some areas, higher detection, but lack systematic evaluation^28^, and according to our evaluation risk conflation with deciduous forests in several areas. Deciduous forest types are generally challenging for both rubber and forest maps, with forest maps prone to omissions and rubber maps prone to conflation. In insular Southeast Asia, all region-wide rubber products assessed here had low detection power, meaning there is a need to expand targeted local mapping approaches^19^. Several older rubber maps^23,25,36,41^ had lower accuracy in our evaluation and have likely been superseded by recent products, including Wang et al.^25^ by the map presented here. Regarding forest maps, recent Joint Research Centre^31,33^ and Google DeepMind/Nature Trace^30,34^ products performed best at differentiating forest and rubber. These are also the only products that map “forest” in exact accordance with the EUDR definition – a definition that in the case of rubber creates a potential area of ambiguity*. However, the best-performing map was a national forest map for Thailand, highlighting the value of local field data-based maps where available. Finally, particularly global and regional-wide products are often not fully independent; reuse of training data or masks derived from other maps create a risk of circular support between maps.

### Rubber-associated clearances

The presence of rubber within forested and protected areas indicates that expansion has often been associated with forest loss, but the scale, timing and land-use context vary strongly across the region. Clearances in Cambodia and Laos were often recent, whereas impacts elsewhere in the region tended to occur earlier. Although this study does not capture post-2016 dynamics, the potential for further clearances in countries with expanding rubber areas may warrant attention. In Cambodia, approximately 1 Mha of concessions may have been granted for rubber^46^, including within forested and protected areas^7^. We mapped rubber in around 20% of the concession area, with large areas still under forest in 2025, indicating future clearance risk.

Clearances in the 2010s were predominantly associated with industrial-scale plantations and/or clusters of plantations, suggesting that concessionary and investment-linked expansion played a larger role than smallholder production.

Our historical estimates bounded the plausible scale of rubber-associated clearance between 1.3-3.0 Mha since 1990. These bounds fall between previously proposed figures: estimates by Hurni and Fox^36^ and Wang et al.^25^ exceed our upper bounds, and the estimate by Sheil et al.^27^ is comparatively low, particularly given it spans a period of rapidly rising rubber prices during which the rubber area in the Mekong region more than doubled^5^ with new plantings potentially exceeding 3 Mha^48^.

### Definitions matter

Estimates of “deforestation” are highly sensitive to definitional and methodological choices. Sheil et al.^27^ used a layer for primary humid forest^43^ to argue that the “actual extent of deforestation” due to rubber plantations in Southeast Asia since 2001 was only 0.6 Mha. If forest is defined narrowly as primary humid forest, this estimate is broadly consistent with our findings. However, rubber, a facultatively deciduous tree, can be cultivated in seasonally water-limited environments and - partly driven by policy (see below) - has expanded into non-humid^49^ and non-primary^6^ forest landscapes.

The definition adopted by Sheil et al.^27^ is useful for crops that primarily replace humid tropical forest, and it has the advantage of low conflation of rubber and agroforest with forest. It is, however, insufficient for a comprehensive region-wide assessment of rubber’s biodiversity impacts. Recent rubber expansion in the Mekong region has often affected drier, degraded and/or regenerating tree cover. Examples include deciduous forests in biodiversity-rich landscapes in the Attapeu plains (Laos), Chư Prông (Vietnam), and Mondulkiri (Cambodia), and secondary forests within customary shifting cultivation systems in northern Laos and northern Myanmar. A focus on primary humid forest therefore only provides a partial assessment of impacts.

This matters because forest types excluded by such a narrow focus have been, and in some places still are, particularly vulnerable to conversion under policies permitting concessions on land classified as “degraded”, “fallow”, “unused” or “unproductive”^11,13,50,51^. Such classifications can be ambiguous and are often contentious^50^. For example, when definitions of “degraded” rely on structural or volumetric thresholds, they may include naturally sparse or regenerating forests^11,13,50^. The notion of “unused” can ignore customary land use^50,51^. While protecting primary forest is essential, and rubber cultivation on genuinely degraded land may entail relatively limited biodiversity loss while supporting livelihoods and carbon sequestration, not all clearances of non-primary tree cover identified here occurred in highly degraded systems. Our findings, together with previous work on social risks of policies permitting industrial tree-plantation concessions on ambiguously classified land^6,11,13,50,51^, suggest a need to monitor impacts in both primary and non-primary forest regions to avoid unintended biodiversity loss and adverse social impacts.

### Context matters

Beyond definitional challenges in estimating “deforestation,” interpretation is highly context-dependent. The ecological and social consequences of smallholder-driven conversion of limited areas differ markedly from large-scale transformations of entire landscapes into rubber monocultures. The sheer scale of conversion of shifting-cultivation landscapes with remnant forest patches across the Mekong region makes negative biodiversity outcomes likely, even where some of the replaced tree cover was already degraded. In addition, reported cases of land dispossession and unequal contract-farming arrangements in Laos^52^, Myanmar^53^ and Cambodia^7^ point to potential social harm, while carbon sequestration benefits remain contested^54^. Remote sensing alone is often insufficient to assess the biodiversity, ecological, and social value of the systems lost and those replacing them, and aggregate deforestation metrics can obscure these complexities.

### Limitations and uncertainties affecting clearance estimates

Our estimates rely on accurate reconstruction of land-cover trajectories from satellite imagery. This was often challenging, particularly for the 1990s-2000s, equatorial areas, and heterogeneous treescapes where semi-natural and planted tree cover form a continuum. Furthermore, extrapolations ignore that rubber-associated clearances are frequently spatially and temporally clustered. Overestimation may have occurred where a small number of clearances were extrapolated across large rubber areas, for example in Indonesia and Thailand. Harder-to-detect clearances may be underestimated, for example in deciduous, sparse or young secondary forests, before the availability of very-high-resolution imagery, and/or in the context of small-scale agriculture. Finally, temporal gaps in imagery mean that attribution was often uncertain. Impacts may be overestimated where areas were initially cleared for another purpose, and underestimated where areas cleared for rubber were left unproductive, later converted to crops such as oil palm, or where rubber indirectly displaced other crops into forests. Further sources of uncertainty and limitations are described in the Methods.

### From mapping to policy

While “deforestation” estimates are context- and definition-dependent, evidence of environmental and social risks supports stronger due diligence in rubber supply chains. At the same time, satellite-mapping limitations and documentation gaps require careful consideration: absence of map-based confirmation or supporting documentation does not necessarily indicate deforestation risk. Producers and operators may face technical limits in demonstrating compliance, especially in equatorial areas and smallholder systems.

Particular caution may be needed when applying overly rigid mapping and traceability requirements to smallholder supply chains. Rubber is produced by around six million smallholders, whose heterogeneous plantations can be difficult to map, misclassified as forest, or intersect with forest maps because of geolocation uncertainty and complex boundaries. When rubber prices are low, temporarily untapped smallholder plantations may also develop naturally regenerating vegetation, complicating binary forest/non-forest classifications. Resulting deforestation alerts are difficult to verify across dynamic supply chains in which rubber often passes through multiple informal intermediaries. Smallholder cooperatives equipped with digital traceability systems^55^ remain the exception rather than the norm. This could make smallholder supply chains appear disproportionately burdensome^†^ or risky and favour more easily documented industrial production.

Such a shift may be counterproductive given rubber’s livelihood importance and our finding that recent clearances were often associated with larger-scale plantations. Industrially produced rubber can move through equally complex cross-border supply chains and connect to higher-risk geographies. For example, Vietnam is a major investor in Cambodian rubber expansion and imports large volumes from Cambodia, while China is Vietnam’s main export market^56^. Although the EU imports little rubber directly from Cambodia, Vietnam is among its top suppliers of natural rubber, and China is a major source of tyre imports. These trade linkages are not well reflected in current EUDR country-level benchmarking: Vietnam, Laos and China are classified as low risk under the EUDR, and Cambodia as standard risk. Tracing these linkages is challenging, suggesting that smallholder production may not be the main source of traceability or deforestation risk.

Effective policy implementation could therefore combine support for smallholders with targeted scrutiny of industrial, concession-linked and investment-linked rubber expansion. This will require cooperation between producer and consumer countries and dialogue among policymakers, stakeholders and scientists to achieve environmental and social goals without marginalising smallholders or encouraging substitution with more carbon-intensive synthetic rubber.

## Methods

We mapped rubber plantations across Southeast Asia using a random forest classifier, integrating optical and radar satellite imagery, phenological metrics (adapted from Wang et al.^25^), canopy height, and spatial texture variables. The mapping was done in Google Earth Engine^57^. To estimate the scale of conversion of natural and semi-natural tree cover associated with rubber expansion, we primarily relied on visual interpretation of historical satellite imagery. This was because map-based and algorithmic separation of planted and natural tree cover is challenging across Southeast Asia, where forests exhibit substantial spectral and structural heterogeneity and the availability of historical forest maps is limited. Detailed methods, and associated uncertainties and limitations are described below.

### Reference data

We compiled a total of 12,449 reference sample points (Fig. S4): 3,578 for rubber, 7,094 for forest (including deciduous, evergreen, degraded, and primary forest), and 1,777 points for other tree plantations (for example, *Acacia* spp., *Eucalyptus* spp., cashew, mango, oil palm, and teak). Of these 3,826 points were sourced from Wang et al.^25^ (over 50% of which are field observations) and the remainder generated through visual interpretation of very high-resolution imagery in Google Earth Pro^58^. Where available the interpretation was supported by Google Street View, information shared by rubber supply chain actors, publicly available information on plantations, and time series imagery from Sentinel-2 and Landsat 5, 7, and 8.

For map validation we generated an additional 2,500 points using the *stratifiedSample* function in Google Earth Engine (Fig. S5). Where available, interpretation was supported by other information as above. Validation points were generally assessed by at least two, and in cases of uncertain interpretation, three interpreters. Points with an uncertain and/or conflicting interpretation were flagged for subsequent sensitivity analyses (see below).

### Rubber phenology, study area and analysis years

Rubber plantations exhibit a characteristic phenological cycle in which trees older than four years undergo brief defoliation during the dry season, followed by rapid refoliation^59^. The timing and duration of defoliation are influenced by macro- and micro-climatic conditions, as well as genotype-environment interactions, with different rubber clones often exhibiting distinct wintering behaviour under the same climatic conditions. Despite some variability, rubber defoliation is generally associated with seasonal water stress, i.e., it broadly coincides with the dry season, and the short leaf-off period (typically 10–15 days) provides a key spectral feature for distinguishing rubber from other tree cover.

In mainland and northern insular Southeast Asia, the dry season typically occurs between January and February, and in equatorial insular Southeast Asia between June and September (Fig. S6). These differences reflect regional monsoon dynamics and the seasonal migration of the Intertropical Convergence Zone^60^. Following the approach of Wang et al.^25^, we therefore divided the study area into two climatic zones: zone A (all of Cambodia, Laos, Myanmar, Thailand, Vietnam, Malaysia, Xishuangbanna and Hainan in China, and parts of Indonesia), where the classification algorithm assumed rubber defoliation to occur between January and February and refoliation to be complete by April; and zone B, comprising areas in Indonesia where multi-source precipitation data^61^ for 2007-2018 indicated that the driest month occurs between June and September (Fig. S6). In zone B, the defoliation was assumed to occur between June and September and refoliation to be complete by December.

While rubber trees tend to be temporarily leafless before rapidly refoliating in regions with a clear-cut dry season, in more humid equatorial climates (southern part of zone A and most of zone B; Fig. S7) leaf abscission and renewal can happen more gradually within individual trees and across stands^59^. This reduces the detectability of rubber using a phenology-based approach, and together with more persistent cloud and haze obscuring optical imagery, partly explains the higher omission errors in Indonesia and parts of Malaysia.

Climatic zone A was mapped using imagery from 2021 because the dry season in the target year 2020 had wetter-than-average conditions linked to the developing 2020–2021 La Niña event. This affected the detectability of rubber phenology (Fig. S8). Climatic zone B was mapped using imagery from 2019, when El Niño-associated dry conditions intensified seasonal drought and defoliation signals there (Fig. S8). Given long rotation times for rubber, the map can be considered broadly representative of plantation distribution in 2020.

### Remote sensing data

#### Optical

For optical data, we used the level 2A Sentinel-2 Surface Reflectance collection obtained through Google Earth Engine^57^. Quality masking to remove pixels affected by cloud and cloud shadow was undertaken using Google’s Cloud Score+. We used annual band medians for the 10 m visible and near-infrared bands and the 20 m red-edge and shortwave infrared bands, and calculated six standard spectral indices: enhanced vegetation index (EVI), normalized difference vegetation index (NDVI), normalized difference water index (NDWI), normalized burn ratio (NBR), modified NBR (MNBR), and soil-adjusted vegetation index (SAVI).

The Sentinel-2 collection was also used to calculate phenology metrics. Specifically, we calculated the 15th percentile of both NBR and NDVI within the assumed defoliation window, their 85th percentile within the assumed refoliation window, and the difference between these percentiles (= magnitude of defoliation). We also calculated defoliation duration using a one-month moving average of NBR from two months prior to the onset of defoliation to two months following the end of refoliation. Defoliation timing and duration were estimated as the period during which NBR values were below the NBR time series mean.

#### Radar

We used both annual time series of 10 m C-band Sentinel-1 Ground Range Detected imagery and the global 25 m L-band PALSAR-2 annual mosaic. For Sentinel-1, Interferometric Wide Swath acquisitions from descending orbits were used, except for Hainan, where descending-pass imagery was unavailable. Annual summary statistics were derived from the VH cross-polarised Sentinel-1 time series (5th, 15th, 50th, 85th, and 95th percentiles) to characterise central tendencies and seasonal variation in radar backscatter (e.g. associated with vegetation phenology). From the PALSAR-2 annual mosaic, we used the HV cross-polarised band.

#### Canopy height and texture

Because plantation canopies tend to exhibit a more homogeneous spatial texture than natural forests we also calculated the standard deviation of both the median annual NBR and of a 2020 canopy height map^62^ within 20, 50, and 100 m neighbourhoods surrounding each pixel.

To account for differences in characteristics such as the onset and duration of defoliation across climatic zones, latitude and longitude were also included in the classifier. For an overview of all input variables see Tables S14-S15 and for correlations between them Figure S9.

### Random Forest Classifier

Classification was performed in Google Earth Engine using a random forest model. Hyperparameters were tuned using out-of-bag error, resulting in a model with 120 trees (130 trees in climatic zone B) and 7 variables per split. Due to computational constraints within Earth Engine, the study region was divided into tiles for model prediction.

In some areas, persistent cloud cover during the defoliation and refoliation periods prevented calculation of the phenology variables. To address this, a second random forest model excluding phenology variables was trained and used to gap-fill these regions (affecting small areas in Sumatra, northern Laos and the north of Myanmar). Areas with persistently unavailable optical imagery were assigned a no-data value in the final map (1% of the study area).

### Masks

A series of masks was applied to the rubber classification output to remove areas of potential conflation. The applied masks were relatively strict, partly explaining the conservative nature of the map.

Three masks were derived from the Sentinel-2 NBR time series: First, areas with an annual median NBR above or equal to 0.65 were excluded to reduce confusion with other denser tree plantations such as *Acacia* spp. Second, areas where minimum NBR during the defoliation window did not fall below 0.5 were also excluded to reduce conflation with systems that show some dry-season defoliation but where this is weaker or more gradual than in rubber (e.g., some deciduous forests, teak and *Acacia* plantations). Third, areas where the median NBR was below 0.4 during the two months before the defoliation window or two months after the refoliation window were masked, as a low NBR outside of the rubber defoliation window is more typical of clearances or more gradual refoliation. Gradual refolation extending into the rainy season is often observed in deciduous forests, however, some species dominant in deciduous forests, such as *Shorea obtusa*, can exhibit a similar phenology to rubber with a short and entirely leafless period of less than 3 weeks followed by rapid leaf flushing occurring before the onset of the rainy season^63^. Consequently, further forest deciduous masks were needed (below).

We applied four additional natural forest masks: First, all areas mapped as forest in 2020 according to a national forest map for Thailand^39^ were excluded. In all other countries, areas with a probability ≥0.84 of being natural forest according to the Natural Forests of the World dataset^30^ were masked out. To remove any remnant confusion with evergreen forest, we masked pixels mapped as primary humid forest^43^ in 2001 that had not experienced tree cover loss between 2001 and 2021 according to Hansen et al.^64^. Finally, the Tropical Moist Forests transition layer^65^ was used to exclude areas classified as undisturbed forest, or disturbed/converted where these changes occurred after 2021. Our analyses of these forest layers (Table S8) suggest that they only rarely conflate rubber with natural forest, particularly in continental Southeast Asia. However, some conflation occurs in insular Southeast Asia, meaning that the masks may have removed some genuine rubber there.

Finally, we removed all pixels mapped as oil palm^66^ and coconut^67^ in other products that have achieved a high-accuracy. To minimise classification noise and artifacts at the pixel level, we applied a focal filter to the classified image (5 × 5 pixel majority filter) and applied a minimum patch size threshold of 0.5 ha to remove isolated pixels. We used ESA WorldCover^68^ to subset the map to areas mapped as tree cover in 2020.

### Accuracy assessment and area estimation

Accuracy and area estimation followed approaches recommended by Olofsson et al.^37^ and were implemented using the R package *mapaccuracy*^69,70^. In the assessment, we distinguished two classes: rubber and other tree cover. Validation samples were allocated using a compromise between proportional and equal allocation: higher sampling intensity was assigned to rubber to robustly estimate commission errors, while some class-area weighting was retained to support stable area estimation. In total, 600 validation points were allocated to rubber and 1,900 to other tree cover.

We did not include a third “non-tree cover” class because this domain is spatially extensive and highly heterogeneous, encompassing large areas where rubber occurrence is highly unlikely, including urban areas, water bodies, barren land, and cropland. Unless sampled extensively, inclusion of this class - argued by some alternative approaches^27^ to be necessary - could disproportionately influence area-adjusted estimates. We therefore focused the validation on rubber versus other tree cover, where the most plausible classification errors occur.

To account for interpreter variability, we additionally evaluated the robustness of the results under a conservative assumption in which all instances of interpreter disagreement and/or uncertainty were treated as map errors (Tables S1, S3–S4).

### Estimating post-rubber boom clearances (2011-2016)

Estimating past clearances for rubber is challenging as most historical “forest” maps do not - or only imperfectly - distinguish rubber from natural forest owing to spectral similarity. As a first step, we evaluated the utility of several maps - including widely used land cover products^71–73^ - for quantifying forest to rubber conversions. This was done by visually inspecting the land cover trajectory at randomly sampled points in areas mapped as rubber by our study and “forest” by these products (200 points per product; 1,200 points in total). Of the evaluated maps, only two exhibited comparatively low rubber–forest conflation rates: (1) a primary humid forest layer for 2001^43^ and (2) a deciduous dipterocarp forest layer^44^ for 2011.

To approximate the distribution of primary humid forest in 2011 we removed from the 2001 primary humid forest layer all pixels that experienced tree cover loss between 2001 and 2010 according to Hansen et al.^64^. A small area that overlapped with the dry dipterocarp forest layer was also removed to avoid double counting (2 kha where we map rubber in ∼2020).

Because these two forest layers do not capture the full extent of regional forest cover, we additionally estimated potential loss of “other” forest by identifying areas outside these layers where the LandTrendr algorithm^45^ detected tree cover disturbance between 2011 and 2016. Specifically, we used the Landsat-based LandTrendr temporal segmentation algorithm to identify pixels where the first recorded disturbance event since 1993 occurred between 2011 and 2016, and where pre-disturbance NBR exceeded 0.6.

We then quantified the spatial overlap between the 2020 rubber map and the three forest layers, i.e. areas where forest to rubber transitions may have occurred. To account for false forest positives (for example long-rotation plantations incorrectly identified as natural forest), we randomly sampled 600 points within the mapped intersections for visual interpretation using very high-resolution imagery in Google Earth Pro (200 points for each layer). The observed false-positive rates were then used to downward adjust estimated forest-to-rubber conversions on a country-by-country and layer-by-layer basis. We did not account for false negatives in the forest layers. False negatives may occur particularly in younger secondary forests and other naturally sparse systems, because the NBR threshold of 0.6 used in the LandTrendr algorithm restricted the analysis to areas with relatively dense initial tree cover.

### Approximating the scale of historical clearances (1990-2016)

To approximate the scale of historical losses of natural and semi-natural tree cover associated with rubber expansion, we identified the land cover preceding rubber establishment for 586 randomly sampled rubber points (derived from the accuracy assessment dataset above). Interpretation of land cover trajectories was conducted primarily using historical imagery available in Google Earth Pro, supplemented with Sentinel-2 and Landsat 5, 7, and 8 time-series imagery accessed through Collect Earth^74^. Where available, interpretation was additionally supported by Google Street View, information shared by rubber supply-chain actors, and publicly available information.

Specifically, interpreters classified prior land cover into four categories: already modified, unknown, semi-natural tree cover, and natural tree cover (Table S17). Potential loss of natural and semi-natural tree cover was then estimated by extrapolation, scaling the proportion of sampled rubber points associated with prior natural or semi-natural tree cover to the total mapped rubber area according to official statistics and our estimated rubber areas.

The above approach has advantages and limitations. A limitation is that extrapolation from sample points does not account for the spatially and temporally heterogeneous nature of clearances, having occurred at different times and under different policy and market contexts across countries. In addition, interpreter assessments are necessarily subject to some degree of subjectivity. Consequently, our results are best interpreted as bounded rather than point estimates. To understand where and when interpretations were most uncertain, interpreters were asked to assign confidence scores ranging from 0.75 (uncertain) to 1 (virtually certain). An advantage of our approach is that it avoids limitations associated with algorithmic separation of planted and natural tree cover. Natural forests in the study area exhibit substantial spectral heterogeneity, with spectral indices such as NBR ranging from well below to well above the values observed in rubber and other plantations – and historical forest maps are largely limited to primary evergreen forests. This is insufficiently representative of forest cover in the region, in particular in continental Southeast Asia (Table S8). Consequently, purely map- or algorithm-based approaches may either overestimate or underestimate forest conversion associated with rubber expansion.

Over 90% of points were independently assessed by at least two interpreters, in some cases three. Points with differing interpretations were subsequently reviewed by all interpreters and, if disagreement was not resolved, assigned to the “unknown” category. To account for the possibility that points with unknown trajectory obscure additional losses, and to assess the robustness of our estimates to interpreter variability, we repeated the analyses by assigning, for each point, the least and most conservative interpretation across interpreters, respectively, along the gradient: modified land → unknown → semi-natural tree cover → natural tree cover.

### Limitations and uncertainties in clearances estimates

The following sources of uncertainty may have contributed to both over- and under-estimation of clearances:

1. **Interpretation challenges:** Our analysis relied on accurate interpretation of land cover trajectories at reference points. However, establishing the naturalness and condition of tree cover is inherently difficult from satellite imagery alone. Limitations affecting rubber mapping - poor image quality in equatorial regions and for earlier time periods, and ambiguity in heterogeneous treescapes where planted and natural tree cover grade into each other – also apply here. Uncertainty was greatest in Indonesia but also affected other regions where distinctions between planted (e.g. orchards, woodlots) and (semi-)natural tree cover were difficult to establish.
2. **Sampling and extrapolation uncertainty:** Estimates were extrapolated from random sample points, but deforestation is spatially and temporally clustered and clearances range from very small- to very large-scale. Consequently, overestimation may have occurred where few smaller clearance events were extrapolated across extensive areas (for example Thailand and Indonesia), or, where, following a few earlier clearances in the 1990s, no further clearances occurred. Underestimates may have occurred where clearances affected large contiguous areas insufficiently captured by the sample (for example Cambodia). The extrapolation-based clearance estimates, thus require caution, in particular at the level of individual countries.
3. **Systematic detection biases:** Clearances affecting deciduous, naturally sparse and young secondary forests, or preceding very high-resolution imagery, and/or small-scale clearances, are likely underrepresented.
4. **Temporal gaps in imagery:** Intermediate land-cover transitions may have been missed. In some cases, rubber may not have driven the initial clearance.
5. **Attribution uncertainty:** In some cases initial clearances may have been for other crops. For example, more recent clearances in Indonesia may be motivated by oil palm with rubber only planted in margins not suitable for oil palm, leading to overestimation of clearances for rubber. *Vice versa*, clearances may have initially occurred for rubber with areas subsequently converted to oil palm, leading to an underestimation.
6. **Displacement effects:** In some cases rubber may have replaced existing agriculture, with delayed returns and/or low prices driving a need for agricultural expansion elsewhere. Such indirect displacement effects are not captured in our estimates.

## Data availability

All satellite imagery and ancillary spatial datasets used in this study are publicly available from the original data providers. The rubber plantation map generated for this study is available via Zenodo at https://doi.org/10.5281/zenodo.21439869. It is also available as a Google Earth Engine asset **`(projects/rubber-499107/assets/rubber_SEA_2020)`** and can be accessed at: https://code.earthengine.google.com/?asset=projects/rubber-499107/assets/rubber_SEA_2020

## Acknowledgements

This work was funded by the Natural Environment Research Council (NE/X016285/1 and NE/Z503368/1). The Royal Botanic Garden Edinburgh is supported by the Scottish Government’s Rural and Environment Science and Analytical Services Division. We thank M. Loyen, S. Savi, S. Nyquist, O. Cupit, A. Davey and C. Ellis for comments on the manuscript.

## Author contributions

A.A. and J.G. conceived the study with input from J.H. and P.H. S.H. developed the computational framework for the rubber map, and A.A. led subsequent analyses. J.H., Y.W. and J.X. contributed data and supported interpretation and analysis. A.A. wrote the manuscript with input from all authors.

## Competing interests

The authors declare no competing interests.

## Inclusion and ethics

This work resulted from a collaborative partnership between scientists and rubber supply-chain actors within and outside rubber-growing regions. Consideration was given to citation diversity. The study was approved by the institutional ethics committee of the Royal Botanic Garden Edinburgh.

## Additional information

Supplementary Information is available for this paper.

Correspondence and requests for materials should be addressed to Antje Ahrends or Sam Harrison.

## Footnotes

* The EUDR definition of forest closely follows the FAO definition. However, plantations of relevant agricultural commodities, including rubber cultivated for latex, are treated as agricultural use and are therefore excluded from the definition of forest, whereas plantations established for wood production may still qualify as forest. This could imply that converting an *Acacia* or *Eucalyptus* plantation to rubber for latex production constitutes deforestation. Further ambiguity arises because rubber plantations may produce latex, wood, or both. Similar land-use changes could therefore receive different regulatory treatment depending on the intended product: conversion to latex- producing rubber may be classified as deforestation, whereas conversion to rubberwood may instead be assessed under the wood provisions, which prohibit the conversion of primary or naturally regenerating forest, but not of planted or plantation forest. Conversely, replacing rubber with a wood plantation could be interpreted as creating forest, such that a subsequent conversion back to rubber might constitute deforestation. These partly paradoxical borderline cases highlight the need for clearer guidance on how conversions between rubber and other plantation forests should be treated.

† To illustrate the scale of manual verification that can be required, in a rubber-trader dataset covering both smallholder and industrial production polygons, 9,114 of 71,825 polygons (13%) overlapped with the JRC Global Forest Cover 2020 v3 map (JRC3); 6,045 polygons (8%) had >10% overlap. Of overlapping polygons, 2,292 had post-2020 tree-cover loss flagged by the Hansen et al. (2013) annual loss product, which in rubber areas can be triggered, for example, by replanting, storm damage, disease or image artefacts. If near-real-time alert products are used, including GLAD-L, GLAD-S2, RADD or DIST-ALERT, false deforestation positives may increase further, as rubber wintering can be mistaken for tree loss. Overlap between these supplier-mapped rubber polygons and JRC3 was most frequent in humid tropical countries, including Cameroon, Malaysia and Indonesia, where the rubber maps assessed here also tended to perform more poorly.

## Notes

https://doi.org/10.5281/zenodo.21439869

https://code.earthengine.google.com/?asset=projects/rubber-499107/assets/rubber_SEA_2020

### Competing Interest Statement

The authors have declared no competing interest.

