## SupplementaryMaterial for "Rubber mapping reveals opportunities and current limitations of deforestation due-diligence"

Draft - not peer reviewed; subject to revision. Findings are preliminary.

### Supporting tables

**Table S1 | Overall accuracy and area estimates.** Calculations follow Olofsson et al.[1] and were implemented using the R package mapaccuracy[2,3]. Around 1% of the study area could not be mapped due to persistent cloud cover. Recalculating accuracy under a conservative assumption, in which all instances of uncertain interpretation (initial interpreter disagreement and/or poor imagery) are treated as map errors, reduced the area-adjusted producer's accuracy to 0.42 and user's accuracy to 0.87. See Table S5 for further details on commission errors.

Validation and area estimates presented in Tables S1, S3 and S4 refer to areas where rubber is an obvious and dominant component of the canopy (generally monocultures). They are likely to underestimate false negative rates, and therefore area, in regions where agroforests are widespread, particularly Indonesia (see Table S9).

**A Error matrix of sample data**

| | | **Reference** | | | |
|---|---|---|---|---|---|
| | | notRubber | rubber | **Total** | **User's accuracy** |
| **Map** | *notRubber* | 1,870 | 30 | 1,900 | 0.98 |
| | *rubber* | 44 | 556 | 600 | 0.93 |
| | Total | 1,914 | 586 | 2,500 | |
| | Producer's accuracy | 0.98 | 0.95 | | |
| | F1 score | 0.98 | 0.94 | | |

**B Error matrix of area proportions**

| | | **Reference** | | | | |
|---|---|---|---|---|---|---|
| | | notRubber | rubber | **Total** | **User's accuracy** | **95% CI** |
| Map | *notRubber* | 0.956 | 0.015 | 0.972 | 0.98 | 0.01 |
| | *rubber* | 0.002 | 0.026 | 0.028 | **0.93** | **0.02** |
| | Total | 0.958 | 0.042 | 1 | | |
| | Producer's accuracy | 1 | **0.63** | **Overall** | **0.98** | **0.01** |
| | 95% CI | 0 | **0.08** | | | |

**C Mapped and estimated area**

| | **Mapped area (ha)** | **Estimated area (ha)** | **Lower 95% CI** | **Upper 95% CI** |
|---|---|---|---|---|
| notRubber | 291,875,182 | 287,893,157 | 286,246,987 | 289,539,327 |
| rubber | 8,543,604 | 12,525,629 | 10,879,459 | 14,171,799 |
| **Total** | 300,418,786 | | | |

Draft - not peer reviewed; subject to revision. Findings are preliminary.

**Table S2 | Estimates of rubber area by country.** Individual country estimates should be treated with caution due to the low number of data points for some countries. This is reflected in wide confidence intervals.

**Column key:** UA = user's accuracy; PA = area-adjusted producer's accuracy; OA = overall accuracy; CI = 95% confidence interval

| Country | N points | UA | CI (UA) | PA | CI (PA) | OA | CI (OA) | Mapped area (ha) | Estimated area (ha) | CI (estim. area) |
|---|---|---|---|---|---|---|---|---|---|---|
| Cambodia | 79 | 0.96 | 0.07 | 1.00 | 0.00 | 1.00 | 0.00 | 391,588 | 377,603 | 27,411 |
| China | 88 | 1.00 | 0.00 | 0.76 | 0.25 | 0.92 | 0.10 | 951,590 | 1,255,724 | 410,259 |
| Indonesia | 1,076 | 0.95 | 0.05 | 0.31 | 0.10 | 0.98 | 0.01 | 1,200,286 | 3,704,766 | 1,208,728 |
| Laos | 127 | 1.00 | 0.00 | 0.59 | 0.47 | 0.99 | 0.02 | 228,041 | 384,910 | 307,457 |
| Malaysia | 241 | 0.73 | 0.11 | 0.67 | 0.31 | 0.98 | 0.02 | 866,646 | 948,724 | 443,745 |
| Myanmar | 314 | 0.95 | 0.06 | 0.65 | 0.31 | 0.99 | 0.01 | 620,614 | 906,651 | 437,405 |
| Thailand | 392 | 0.95 | 0.03 | 0.81 | 0.13 | 0.96 | 0.03 | 3,435,939 | 4,040,475 | 676,635 |
| Vietnam | 183 | 0.85 | 0.09 | 0.82 | 0.29 | 0.99 | 0.02 | 848,900 | 878,460 | 321,482 |
| Total area | 2,500 | 0.93 | 0.02 | 0.63 | 0.08 | 0.98 | 0.01 | 8,543,604 | 12,525,629 | 1,646,170 |

Draft - not peer reviewed; subject to revision. Findings are preliminary.

**Table S3 | Accuracy and area estimates for continental Southeast Asia.** Here we use “continental Southeast Asia” to refer to Myanmar, Thailand, Cambodia, Laos, Vietnam, Xishuangbanna and Hainan (i.e. excluding peninsula Malaysia). Recalculating accuracy under a conservative assumption, in which all instances of uncertain interpretation are treated as map errors, reduced the area-adjusted producer’s accuracy to 0.65, while user’s accuracy remained high (0.93). Calculations were performed as described in Table S1.

**A Error matrix of reference data**

| | | **Reference** | | | |
|---|---|---|---|---|---|
| | | notRubber | rubber | **Total** | **User's accuracy** |
| **Map** | *notRubber* | 717 | 11 | 728 | 0.98 |
| | *rubber* | 24 | 431 | 455 | 0.95 |
| | Total | 741 | 442 | 1,183 | |
| | Producer's accuracy | 0.97 | 0.98 | | |
| | F1 score | 0.98 | 0.96 | | |

**B Error matrix of area proportions**

| | | **Reference** | | | | |
|---|---|---|---|---|---|---|
| | | notRubber | rubber | **Total** | **User's accuracy** | **95% CI** |
| Map | *notRubber* | 0.932 | 0.014 | 0.946 | 0.98 | 0.01 |
| | *rubber* | 0.003 | 0.051 | 0.054 | **0.95** | **0.02** |
| | Total | 0.935 | 0.065 | 1 | | |
| | Producer's accuracy | 1 | **0.78** | **Overall** | **0.98** | **0.01** |
| | 95% CI | 0 | **0.1** | | | |

**C Mapped and estimated area**

| | **Mapped area (ha)** | **Estimated area (ha)** | **Lower 95% CI** | **Upper 95% CI** |
|---|---|---|---|---|
| notRubber | 114,242,864 | 112,858,294 | 111,836,520 | 113,880,068 |
| rubber | 6,476,672 | 7,861,242 | 6,839,468 | 8,883,016 |
| **Total** | 120,719,536 | | | |

**Table S4 |** Accuracy and area estimates for insular Southeast Asia. Here we use the term ‘insular Southeast Asia’ to refer to all of Malaysia (including Peninsular Malaysia) and Indonesia. Calculations were performed as described in Table S1.

The below estimates apply to plantations where rubber constitutes a dominant component of the canopy and is detectable in the very high-resolution imagery used for validation. Recalculating accuracy under a conservative assumption, in which all instances of uncertain interpretation were treated as map errors reduced the area-adjusted producer’s accuracy to 0.16 and the user’s accuracy to 0.68. Cases of uncertain interpretation often affected suspected rubber agroforests with mixed canopies, which tend not to be captured by our approach (see also Table S9).

**A Error matrix of reference data**

| | | **Reference** | | | |
|---|---|---|---|---|---|
| | | notRubber | rubber | **Total** | **User's accuracy** |
| **Map** | *notRubber* | 1,153 | 19 | 1,172 | 0.98 |
| | *rubber* | 20 | 125 | 145 | 0.86 |
| | Total | 1,173 | 144 | 1,317 | |
| | Producer's accuracy | 0.98 | 0.87 | | |
| | F1 score | 0.98 | 0.87 | | |

**B Error matrix of area proportions**

| | | **Reference** | | | | |
|---|---|---|---|---|---|---|
| | | notRubber | rubber | **Total** | **User's accuracy** | **95% CI** |
| Map | *notRubber* | 0.972 | 0.016 | 0.988 | 0.98 | 0.01 |
| | *rubber* | 0.002 | 0.01 | 0.012 | **0.86** | **0.06** |
| | Total | 0.974 | 0.026 | 1 | | |
| | Producer's accuracy | 1 | **0.38** | **Overall** | **0.98** | **0.01** |
| | 95% CI | 0 | **0.11** | | | |

**C Mapped and estimated area**

| | **Mapped area (ha)** | **Estimated area (ha)** | **Lower 95% CI** | **Upper 95% CI** |
|---|---|---|---|---|
| notRubber | 177,632,318 | 175,037,707 | 173,747,585 | 176,327,829 |
| rubber | 2,066,932 | 4,661,543 | 3,371,421 | 5,951,665 |
| **Total** | 179,699,250 | | | |

**Table S5 | List of rubber map commission errors detected during map validation.** In total 44 of 600 points mapped as rubber were false positives. Misclassifications occurred for example in tree plantations, open forest types or clearings, when clearing, leaf loss or browning during the dry season followed by rapid greening mimicked the phenological signal of rubber. Additional commission errors resulted from the use of focal mean filters (see Methods), causing small patches of non-rubber vegetation embedded within rubber stands to be classified as rubber. Most of the commission errors were found in Malaysia (26%), followed by Vietnam (15%). For example, in northern Vietnam confusion was observed with an unidentified broadleaf plantation, and in Malaysia with palm. The reasons for confusion with evergreen palms are unclear but may relate to generally reduced map performance in insular Southeast Asia where palm plantations are widespread.

| Landcover category | Landcover type | False positives (n) |
|---|---|---|
| **Forest** | Deciduous forest | 2 |
| | Degraded forest | 2 |
| | Unknown forest type | 1 |
| **Plantation** | Acacia | 1 |
| | Unknown broadleaf tree plantation | 2 |
| | Palm | 12 |
| | Unknown tree plantation | 5 |
| **Other** | Clearing | 8 |
| | Cropland | 4 |
| | Grass | 1 |
| | Sparse trees | 1 |
| | Flood plain | 2 |
| | Unknown | 3 |

Draft - not peer reviewed; subject to revision. Findings are preliminary.

**Table S6 | Overlap of areas mapped as rubber in this study and areas mapped as forest in global forest datasets.** While overlap may reflect misclassification in either map, cases of overlap affecting more than a million hectares (JRC V1, GFT V0, SBTN V1 and V2) suggest substantial conflation in forest datasets as our rubber-mapping approach was designed to minimise commission errors with natural forest. See also Table S7.

**Column key:** JRC V1-V3 = versions 1-3 of the EC JRC Global Forest Cover 2020 map[4-6]; GFT V0 and V1 = versions 0 and 1 of the EC JRC Global Forest Types 2020 map[7,8] (including classes 'primary forest', 'naturally regenerating forest' and 'Planted/Plantation forest'); GFT V1 is treated as equivalent with JRC V3 as the extent of JRC V3 was used as the base layer for mapping GFT V1; NFW = Natural Forests of the World[9] using the OA probability threshold (0.52) reported by the authors; ForTy = Global Forest Typology[10] (including classes 'primary forest', 'naturally regenerating forest', 'plantation forest' and 'planted forest'); SBTN = versions 1 and 2 of the Science-Based Targets Network Natural Lands map[11] (including classes 'forests', 'mangroves', 'wet forests', and 'peat forests').

Note that the forest layers differ in their definition of "forest": JRC, GFT and ForTy include planted and plantation forests to align with EUDR definitions, whereas NFW and SBTN map only natural forest areas.

| | JRC V1 | JRC V2 | JRC V3 / GFT V1 | GFT V0 | NFW | ForTy | SBTN V1 | SBTN V2 |
|---|---|---|---|---|---|---|---|---|
| Cambodia | 87,056 | 14,831 | 6,406 | 43,979 | 8,686 | 11,854 | 70,276 | 70,341 |
| China | 920,218 | 71,990 | 89,789 | 77,692 | 5,573 | 3,870 | 272,301 | 273,329 |
| Indonesia | 290,655 | 149,569 | 41,224 | 212,495 | 80,470 | 59,517 | 11,684 | 21,989 |
| Laos | 139,957 | 14,587 | 11,136 | 68,399 | 18,051 | 8,307 | 126,611 | 126,288 |
| Malaysia | 68,186 | 34,003 | 20,184 | 50,023 | 31,573 | 33,371 | 22,675 | 23,239 |
| Myanmar | 435,206 | 96,862 | 32,942 | 100,235 | 24,340 | 13,645 | 166,604 | 166,276 |
| Thailand | 2,196,146 | 301,023 | 70,549 | 695,957 | 92,992 | 38,302 | 416,510 | 416,009 |
| Vietnam | 537,153 | 95,004 | 76,848 | 169,993 | 46,062 | 52,994 | 408,558 | 409,173 |
| **Total** | **4,674,578** | **777,868** | **349,079** | **1,418,772** | **307,747** | **221,860** | **1,495,218** | **1,506,642** |

Draft - not peer reviewed; subject to revision. Findings are preliminary.

**Table S7 | Comparison of forest omission and rubber misclassification rates across forest datasets in the study area.** Values are means and the 2.5th and 97.5th percentiles across 100 repeated subsampling iterations (see below).

Forest datasets partly had high rates of conflation with rubber. The best balance was achieved by a national forest map for Thailand[12] (the only official country-level product available to us). In terms of region wide products, recent versions of the JRC Forest Cover[5] and Forest Type[8] 2020 maps, and the Google DeepMind/WRI Natural Forests of the World[9] and Forest Typology[10] performed best. Almost all forest datasets suffered from higher rubber conflation in insular than in continental Southeast Asia. Conflation with rubber agroforests will be higher than detected here given their similar appearance on satellite imagery to natural forests. This is also why we restricted the assessment of forest omissions to continental Southeast Asia.

In continental Southeast Asia, more recent forest datasets[5,8-10] tended to have omission rates below 5%. Where omissions occurred, they mainly affected deciduous forest ecoregions, including the Central Indochina Dry Forests, Northern Khorat Plateau Moist Deciduous Forests, and Irrawaddy Dry and Moist Deciduous Forests. These areas were also prone to forest conflation in several rubber maps (Table S8) and may hence require heightened attention in due-diligence assessments. Consistent with their narrower scope, products[13,14] focused on moist forest types had higher omission rates, but have the advantage of low conflation with rubber.

The assessment was based on repeated (n=100) stratified random subsampling of forest and rubber points. Subsampling of forest points was by ecoregion[15], with sample sizes proportional to forest area[16] per ecoregion (one point per 100 kha); and of rubber points by subnational administrative areas, with sample sizes proportional to officially reported plantation areas (one point per 10 kha). The points included this study's training and validation data (Figs. S4, S5). Additional points were collected through visual interpretation from very-high resolution imagery to fill gaps for individual ecoregions and/or rubber-producing provinces. We applied minimum-distance spatial thinning to enforce a minimum spacing of 10 km between forest points and 1 km between rubber points. This resulted in 6,958 points available for subsampling (4,274 for forest and 2,684 for rubber). Sources of officially reported rubber areas are listed in Table S13; for Cambodia only national-level statistics were available.

Note that the forest layers differ in their definition of "forest": primary humid forest and tropical moist forest datasets map narrower forest types, whereas JRC forest-cover maps include planted and plantation forests to align with EUDR definitions. JRC Global Forest Types[7], Forest Typology and De Keersmaecker et al.[17] also include planted and plantation forests, but as separate classes, which allowed us to exclude them here for comparability with datasets restricted to natural forest. For SBTN[11] we included the classes 'forests', 'mangroves', 'wet forests', and 'peat forests' and for Gosling et al.[18] 'likely natural' and 'potentially natural'. Classes in the Tropical Moist Forest[13] dataset were 10: 'undisturbed tropical forest'; 20: 'degraded tropical moist forest'; 70: 'savannah, deciduous forest, agriculture, evergreen shrubland, non-vegetated cover and afforestation'. The primary humid forest layer is only available for 2001; we approximated primary forest in 2020 by removing pixels that experienced tree cover loss between 2001 and 2019 according to Hansen et al.[19]. The original primary humid forest layer for 2001 gave similarly high forest omission rates (56%).

| Dataset | Forest not mapped (%) (continental SEA) | Rubber misclassified as forest (%) (continental) | Rubber misclassified as forest (%) (insular) |
|---|---|---|---|
| JRC[4] V1 | 3 (2 - 4) | 65 (63 - 67) | 23 (21 - 25) |
| JRC[6] V2 | 9 (8 - 10) | 5 (4 - 6) | 14 (13 - 16) |
| JRC[5] V3 | 4 (3 - 5) | 3 (2 - 4) | 9 (8 - 10) |
| JRC[7] Global Forest Types V0 | 7 (6 - 8) | 15 (13 - 17) | 16 (15 - 18) |
| JRC[8] Global Forest Types V1 | 4 (4 - 5) | 2 (1 - 3) | 7 (6 - 9) |
| Forest Typology[10] | 2 (1 - 2) | 1 (0 - 2) | 9 (7 - 10) |
| Rui et al.[20] | 14 (13 - 15) | 18 (16 - 19) | 12 (10 - 14) |
| Natural Forests of World[9] @0.52 | 2 (2 - 3) | 2 (1 - 3) | 11 (10 - 13) |
| Natural Forests of World @0.84 | 10 (8 - 11) | 0 (0 - 1) | 3 (3 - 4) |
| GEM[21] | 11 (10 - 12) | 5 (3 - 6) | 23 (21 - 25) |

| Dataset | Forest not mapped (%) (continental SEA) | Rubber misclassified as forest (%) (continental) | Rubber misclassified as forest (%) (insular) |
|---|---|---|---|
| Natural Forest Persistence[22] @0.57 | 1 (1 - 1) | 77 (75 - 79) | 62 (59 - 65) |
| Natural Forest Persistence @0.62 | 1 (1 - 2) | 70 (68 - 72) | 48 (45 - 51) |
| IPCC Forest Types[23] | 14 (13 - 16) | 9 (8 - 10) | 12 (11 - 13) |
| SBTN[11] V0 | 6 (5 - 7) | 22 (20 - 24) | 3 (2 - 4) |
| SBTN V1.1 | 6 (5 - 7) | 22 (20 - 24) | 4 (3 - 6) |
| TMF[13,24] (classes 10, 20, 70) | 15 (14 - 17) | 71 (69 - 72) | 53 (50 - 56) |
| TMF (classes 10, 20) | 57 (55 - 58) | 1 (0 - 1) | 15 (13 - 16) |
| Primary humid forest[14] in 2020 | 57 (56 - 59) | 0 (0 - 1) | 0 (0 - 0) |
| Lesiv et al.[16] | 7 (6 - 8) | 12 (10 - 14) | 32 (30 - 35) |
| De Keersmaecker et al.[17] | 5 (4 - 6) | 14 (12 - 16) | 27 (25 - 29) |
| Gosling et al.[18] | 13 (12 - 14) | 28 (26 - 30) | 16 (15 - 18) |
| Thailand National Forest Map[12] | 1 (0 - 3) | 1 (0 - 1) | - |

**Table S8 | Comparison of rubber omission and forest misclassification rates across rubber datasets in the study area.** Values (A) show percentage of sample rubber points not detected (omission error), and (B) the percentage of sample forest points misclassified as rubber (commission error relative to forest). For details on the sample data see Table S7.

Compared with other pantropical or region-wide datasets, our map had low forest commission rates and the highest (apparent) performance when evaluated against our validation data and those of Sheil et al.[25]. However, note data limitations discussed below. Higher rubber detection in FDP b in continental Southeast Asia appeared to trade off against increased forest conflation. Forest conflation appeared low in De Keersmaecker et al., but this dataset had higher rubber omission. The map by Wang et al. was outperformed by our map in both respects and can be considered superseded. Maps by Xiao et al. and Condro et al. appeared to perform least well and maps by Petersen et al. and Hurni and Fox (for 2014) are now increasingly outdated. The Kalimantan map by Le Maire et al. for agroforests and monocultures had highest rubber detection in its mapped area.

Forest conflation in rubber datasets mainly affected deciduous and semi-evergreen ecoregions. Examples include the Mizoram-Manipur-Kachin Rain Forests in Myanmar (particularly in FDP b) and the Northern Khorat Plateau Moist Deciduous Forests in northeastern Thailand and northern Laos (particularly in FDP b and Xiao et al.). Most maps had some conflation in the Central Indochina Dry Forests, particularly FDP and b, Xiao. et al. and Wang et al. FDP maps also had conflation in some evergreen forest ecoregions, including the South China-Vietnam Subtropical Evergreen Forests and Northern Vietnam Lowland Rain Forests. Estimated forest conflation in the dataset by Le Maire et al. was below 5% and may be lower still; agroforests can visually resemble natural forests on satellite imagery, hence the conflation rate cannot robustly estimated here.

Recent rubber maps by Le Maire et al., FDP and De Keersmaecker et al., together with our map, are largely complementary and can provide converging evidence of rubber, provided above tendencies to conflate forest in some areas are considered. Conflation by the probability-based FDP products can be reduced by increasing local thresholds; given parts of the forest areas listed above had rubber probability values of 0.9 or higher, we recommend strict thresholding in these ecoregions. Given low rubber detection even in newer products across Sumatra and Malaysia, Petersen et al. may still add evidence there (of non-forest, even where plantation types changed).

**Data limitations:** The map comparison included data used to train our classifier and is likely to favour our map. Our validation data were in part collected by the same interpreters, potentially introducing shared interpretation biases. While independent validation data (by Sheil et al.[25]) showed similar patterns of map performance, these data were compiled to evaluate the map of Wang et al.[26] and are not well suited to evaluating other products. Differences in mapping year and resolution across several products further limit comparability. The comparison between our and other maps is therefore most appropriate for Wang et al.[26], who used imagery for the same year and resolution, and a subset of the same training data. Apparent lower performance of some other products requires further checking.

**Column key:** CHN = China, KHM = Cambodia, LAO = Laos, MMR = Myanmar, VNM = Vietnam, THA = Thailand, MYS = Malaysia, IDN = Indonesia; Kal (SCW) = Kalimantan Barat, Tengah and Selatan provinces; **Row key:** FDP = Forest Data Partnership[27,28] probability models a and b for 2020 (thresholded at values that balance precision and recall in FDP evaluations: 0.59/0.44 using withheld test data and 0.23/0.35 using the dataset by Sheil et al.[25]; the latter includes samples for agroforestry); De Keersmaecker et al.[17] = rubber map for 2020; Wang et al.[26] = rubber map for 2021; SDPT2/SDPT3 = Spatial Database of Planted Trees[29] versions 2 and 3, incorporating datasets by Xiao et al.[30], Debonne et al.[31], Petersen et al.[32] (for 2013–2014) and Condro et al.[33] (for 2019). We included cropping systems mixed with rubber to avoid false appearance of low detection. The SDPT has additional data for unknown and other plantation types (not included here), which can still provide evidence that an area is not forest. For Xiao et al.[30] we use the data in SDPT V2. Xiao et al.[30] report somewhat larger rubber areas than mapped in SDPT as, in areas of overlap, newer datasets take precedence in the SDPT.

**A Rubber points undetected (%)**

| Map | CHN | KHM | LAO | MMR | VNM | THA | MYS | IDN | Kal (SCW) |
|---|---|---|---|---|---|---|---|---|---|
| This study | 9<br>(5 - 14) | 4<br>(0 - 9) | 17<br>(7 - 29) | 5<br>(2 - 10) | 9<br>(5 - 13) | 12<br>(11 - 14) | 39<br>(33 - 44) | 55<br>(53 - 58) | 48<br>(40 - 56) |
| FDP b @0.35 | 3<br>(1 - 7) | 2<br>(0 - 5) | 13<br>(5 - 25) | 2<br>(2 - 3) | 5<br>(1 - 7) | 3<br>(2 - 4) | 43<br>(36 - 49) | 92<br>(91 - 92) | 100<br>(100 - 100) |

Draft - not peer reviewed; subject to revision. Findings are preliminary.

| Map | CHN | KHM | LAO | MMR | VNM | THA | MYS | IDN | Kal (SCW) |
|---|---|---|---|---|---|---|---|---|---|
| FDP b @0.44 | 5 (1 - 8) | 3 (0 - 7) | 13 (5 - 25) | 2 (2 - 3) | 5 (2 - 8) | 4 (3 - 5) | 45 (39 - 53) | 93 (92 - 93) | 100 (100 - 100) |
| FDP a @0.23 | 11 (6 - 17) | - | - | - | 11 (6 - 15) | 22 (20 - 23) | 45 (36 - 51) | 45 (42 - 47) | 67 (61 - 73) |
| FDP a @0.59 | 16 (10 - 22) | - | - | - | 17 (12 - 22) | 31 (29 - 33) | 63 (56 - 69) | 58 (55 - 61) | 79 (73 - 85) |
| De Keersmaecker et al. | 44 (35 - 52) | 24 (15 - 32) | 35 (22 - 49) | 20 (13 - 27) | 40 (34 - 47) | 26 (23 - 28) | 65 (57 - 71) | 62 (58 - 64) | 72 (66 - 78) |
| Wang et al. | 5 (2 - 9) | 8 (2 - 12) | 21 (10 - 36) | 17 (10 - 25) | 13 (9 - 19) | 12 (10 - 14) | 47 (39 - 54) | 56 (54 - 59) | 68 (61 - 73) |
| Xiao et al. | - | 99 (98 - 100) | 83 (74 - 90) | 99 (97 - 100) | 96 (93 - 98) | 50 (48 - 52) | - | - | - |
| Petersen et al. | - | 54 (43 - 65) | - | - | - | - | 54 (45 - 61) | 59 (56 - 62) | 73 (66 - 80) |
| Condro et al. | - | - | - | - | - | - | - | 100 (100 - 100) | 100 (100 - 100) |
| SDPT2 | - | 47 (34 - 58) | 83 (74 - 90) | 99 (97 - 100) | 96 (93 - 98) | 50 (48 - 52) | 54 (45 - 61) | 59 (56 - 62) | 73 (66 - 80) |
| SDPT3 | - | 43 (29 - 53) | 95 (85 - 100) | 99 (98 - 100) | 96 (93 - 99) | 54 (51 - 56) | 54 (45 - 61) | 59 (56 - 62) | 73 (66 - 80) |
| Hurni & Fox | - | 40 (30 - 52) | 65 (50 - 80) | - | 43 (38 - 49) | - | - | - | - |
| Le Maire et al. | - | - | - | - | - | - | - | - | 19 (15 - 24) |

### B Forest points misclassified as rubber (%)

| Map | CHN | KHM | LAO | MMR | VNM | THA | MYS | IDN | Kal (SCW) |
|---|---|---|---|---|---|---|---|---|---|
| This study | 0 (0 - 2) | 0 (0 - 0) | 0 (0 - 0) | 1 (0 - 1) | 1 (0 - 1) | 0 (0 - 0) | 0 (0 - 1) | 0 (0 - 0) | 0 (0 - 0) |
| FDP b @0.35 | 3 (0 - 8) | 2 (0 - 4) | 13 (10 - 16) | 6 (5 - 8) | 12 (10 - 15) | 5 (3 - 6) | 0 (0 - 0) | 0 (0 - 0) | 0 (0 - 0) |
| FDP b @0.44 | 3 (0 - 7) | 1 (0 - 3) | 9 (7 - 12) | 5 (4 - 6) | 8 (6 - 10) | 3 (1 - 4) | 0 (0 - 0) | 0 (0 - 0) | 0 (0 - 0) |
| FDP a @0.23 | 6 (2 - 11) | - | - | - | 12 (9 - 15) | 7 (4 - 9) | 1 (0 - 1) | 1 (1 - 2) | 0 (0 - 1) |
| FDP a @0.59 | 3 (0 - 5) | - | - | - | 8 (6 - 10) | 2 (1 - 3) | 1 (0 - 1) | 1 (0 - 1) | 0 (0 - 0) |
| De Keersmaecker et al. | 2 (0 - 3) | 0 (0 - 0) | 2 (1 - 2) | 1 (0 - 1) | 1 (0 - 1) | 1 (0 - 2) | 0 (0 - 0) | 0 (0 - 0) | 0 (0 - 0) |
| Wang et al. | 0 (0 - 2) | 4 (1 - 8) | 2 (1 - 3) | 2 (1 - 2) | 4 (3 - 6) | 9 (6 - 11) | 1 (0 - 2) | 2 (1 - 2) | 2 (0 - 2) |
| Xiao et al. | - | 0 (0 - 0) | 2 (0 - 4) | 1 (0 - 2) | 1 (0 - 1) | 5 (3 - 7) | - | - | - |
| Petersen et al. | - | 1 (0 - 3) | - | - | - | - | 3 (2 - 4) | 1 (0 - 1) | 0 (0 - 0) |
| Condro et al. | - | - | - | - | - | - | - | 0 (0 - 0) | 0 (0 - 0) |
| SDPT2 | - | 1 (0 - 3) | 2 (0 - 4) | 1 (0 - 2) | 1 (0 - 1) | 5 (3 - 7) | 3 (2 - 4) | 1 (0 - 1) | 0 (0 - 0) |
| SDPT3 | - | 1 (0 - 4) | 1 (0 - 2) | 2 (1 - 2) | 1 (0 - 1) | 4 (2 - 5) | 3 (2 - 4) | 1 (0 - 1) | 0 (0 - 0) |
| Hurni & Fox | - | 0 (0 - 0) | 1 (0 - 2) | - | 0 (0 - 0) | - | - | - | - |

Draft - not peer reviewed; subject to revision. Findings are preliminary.

**Table S9 | Comparison of rubber datasets with a reference map[34] for agroforestry and monoculture rubber in 2020 in Kalimantan.** The reference map covers three provinces (Kalimantan Barat, Tengah and Selatan) and achieved an overall accuracy of 87% against field data[34]. We treated this map as a benchmark to compare our and other pantropical and/or region-wide rubber datasets. All products had very high omission relative to the reference map, failing to detect some 90-100% of the reference rubber area. Where our map did detect rubber, agreement with the reference map was high; lower agreement in several other products may indicate higher commission errors. Overall, insular Southeast Asia remains a weakness in rubber mapping. Better maps are required because forest maps also tend to suffer from elevated misclassification rates there. For example, the most recent EC JRC Global Forest Cover 2020 v3 map[5] (in red) classifies half the reference rubber area as forest.

In addition to low map performance for monoculture rubber, agroforestry rubber is largely missed in maps, while it is classified as forest in forest maps. Agroforests span a continuum from rubber integrated within secondary forests to low-intensity plantations and are characterised by mixed-species canopies and multiple vegetation strata (D.L.A. Gaveau, personal communication, 2026). Their omission in monoculture rubber maps is not necessarily a failure: treating these systems as a separate class, distinguished from both natural forest and monoculture rubber, is useful as conversion from agroforestry to monoculture rubber tends to reduce structural complexity and may negatively affect biodiversity.

**Column key:** FDP = Forest Data Partnership[27,28] rubber probability models 2025a/2025b for 2020 (for thresholds see Table S8); GFM = De Keersmaecker et al.[17] rubber mapped for 2020. Wang et al.[26] = rubber mapped for 2021; Petersen et al.[32] = rubber mapped for 2013–2014; Condro et al.[33] = rubber mapped for 2019. GFM has a general agroforest class. If included, it raises the captured agroforestry to 9% but, due to the combined nature of that class, reduces overlap with the reference map to 19%. Combined classes in Petersen et al.[32] for rubber and other commodities were also not included to avoid a false appearance of high commission errors. Overlap with the reference was low regardless, likely due to differences in mapping period, available image quality, and methods: areas by Petersen et al.[32] were manually delineated from imagery for 2013/14 (J. Richter, personal communication, 2026). Low agreement with the map by Wang et al.[26] reflects extensive commission errors in their product for this region[25]; agreement was also low with maps by Condro et al.[33] and FDP model 2025a.

| | This study | FDPb 0.44 | FDPb 0.35 | GFM | FDPa 0.59 | FDPa 0.23 | Petersen et al. | Wang et al. | Condro et al. | Refe-rence | JRC3 |
|---|---|---|---|---|---|---|---|---|---|---|---|
| **Total** rubber area (kha) | 125 | 0 | 0 | 174 | 171 | 420 | 193 | 994 | 166 | 1,830 | |
| Overlap with reference map (ha) | 106 | 0 | 0 | 120 | 73 | 156 | 50 | 174 | 23 | - | 897 |
| Reference area captured (%) | 6 | 0 | 0 | 7 | 4 | 9 | 3 | 10 | 1 | - | |
| **Overlap of mapped with reference (%)** | 85 | 82 | 80 | 69 | 43 | 37 | 26 | 18 | 14 | **-** | **49** |
| **Monoculture** area (kha) | 94 | 0 | 0 | 106 | 43 | 84 | 40 | 116 | 7 | 616 | |
| Reference area captured (%) | 15 | 0 | 0 | 17 | 7 | 14 | 6 | 19 | 1 | - | 13 |
| Share of mapped (%) | 75 | 56 | 51 | 61 | 25 | 20 | 20 | 12 | 4 | - | |
| **Agroforestry** rubber area (kha) | 12 | 0 | 0 | 14 | 31 | 72 | 10 | 58 | 16 | 1,214 | |
| Reference area captured (%) | 1 | 0 | 0 | 1 | 3 | 6 | 1 | 5 | 1 | - | 67 |
| Share of mapped (%) | 10 | 26 | 29 | 8 | 18 | 17 | 5 | 6 | 10 | - | |

**Table S10 | Overlap between areas mapped as rubber (~2020) and areas mapped as natural forest (~2011) in existing forest datasets.**

**Column key:** Area shows the extent of overlap between rubber and natural forest layers. To account for false positives, we validated random samples from that overlap using very high-resolution imagery. Sample size *n* is the number of samples for which land-cover trajectories could be determined (excluding 53 points classified as unknown and 55 false rubber positives), and % is the proportion of samples that experienced rubber-associated loss of (semi-)natural tree cover between 2011-2016.

Only two forest datasets were available, covering dry dipterocarp[35] and primary humid[14] forests. The primary humid forest dataset is for 2001; to avoid including areas cleared prior to 2011, we removed pixels that experienced tree cover loss between 2002-2010 according to Hansen et al.[19]. We additionally applied the LandTrendr algorithm[36] to detect loss of other natural and semi-natural tree cover elsewhere, as the forest two datasets only cover a subset of forest types occurring in the region. Specifically, we identified all pixels that showed spectral stability from 1993 until a first disturbance between 2011-2016. Candidate pixels for potential clearances were further restricted to pixels with a pre-disturbance Normalized Burn Ratio >0.6, selecting for relatively dense forest.

Overlap between areas mapped as rubber in this study and the primary humid forest layer often indicated genuine losses. This is because the low spectral similarity between rubber and these predominantly evergreen forests lead to low conflation rates (see also Table S7). In contrast, the dry dipterocarp forest layer suffered from substantial conflation with rubber, as deciduous forest and rubber can have similar spectral signatures. This led to more false positives, particularly in Thailand, where there are extensive deciduous forest areas and rubber areas were well developed by 2011. The LandTrendr approach cannot distinguish long-rotation plantations from natural forest, leading to high false positive rates in areas where rubber or other long-rotation plantations were already extensive prior to 1993 (e.g., Indonesia, Malaysia, Thailand, China and Vietnam). Corresponding adjustments were made to the estimated clearance areas (Table 2) on a layer-by-layer and country-by-country basis.

| | **Primary humid forest** | | | **Dry dipterocarp forest** | | | **Other tree cover change** | | |
|---|---|---|---|---|---|---|---|---|---|
| | **Area (ha)** | ***n*** | **%** | **Area (ha)** | ***n*** | **%** | **Area (ha)** | ***n*** | **%** |
| Cambodia | 72,026 | 82 | 94 | 9,207 | 19 | 63 | 54,595 | 11 | 82 |
| China | 1,823 | 2 | 50 | | | | 57,219 | 15 | 0 |
| Indonesia | 21,439 | 16 | 88 | | | | 123,632 | 31 | 13 |
| Laos | 12,092 | 12 | 100 | 3,903 | 8 | 25 | 18,665 | 5 | 40 |
| Malaysia | 30,935 | 20 | 95 | | | | 154,687 | 29 | 3 |
| Myanmar | 7,029 | 6 | 67 | 19,457 | 39 | 62 | 42,767 | 10 | 70 |
| Thailand | 5,662 | 5 | 60 | 25,948 | 49 | 4 | 228,982 | 57 | 5 |
| Vietnam | 22,721 | 25 | 100 | 12,851 | 29 | 31 | 82,089 | 22 | 14 |
| | **173,727** | **168** | **82** | **71,366** | **144** | **37** | **762,636** | **180** | **28** |

**Table S11 | Approximated scale of loss of natural and semi-natural tree cover associated with rubber between 1990 and 2016 by country.** The figures in this table are best interpreted as plausible bounds, with large uncertainty, rather than point estimates with precise confidence intervals.

Sample size *n* denotes the number of points for which prior land cover was interpreted. Estimates of potential clearance are based on the proportion of rubber points where the pre-rubber land cover was classified as natural or semi-natural tree cover. (A detailed description of land cover classes is in Table S17.) These proportions were extrapolated to the total estimated rubber area (per country) using estimates from this study and official statistics. Estimated areas for "unknown" land-cover transitions are based on points that interpreters felt unable to classify (i.e., points classed as unknown or points without consensus). We treated these as a separate class rather than removing them from the denominator used to derive transition proportions, which was more conservative and appropriate given the large size of this class.

Most points were assessed by two, and in some cases three, independent interpreters. Points with disagreement were re-assessed (often with the help of a third interpreter). Ranges (shown in italic grey text) were derived by taking the most and least conservative interpretations across interpreters in cases of initial disagreement.

Interpreter variability was particularly large for Malaysia and Indonesia due to poor image quality and difficult-to-interpret heterogenous tree covers. Estimates for these two countries therefore require particular caution but caution is also needed for other country estimates as the points with interpretable land cover trajectories was often low. This caveat particularly affects countries where a low number of sampled clearances were extrapolated across large rubber areas (e.g. Thailand and Indonesia).

| | | | | **Extrapolating this study (ha)** | | **Extrapolating official statistics (ha)** | |
|---|---|---|---|---|---|---|---|
| **Country** | ***n*** | **Proportion (semi-) natural** | **Proportion unknown** | **(Semi-) natural** | **Unknown** | **(Semi-) natural** | **Unknown** |
| Cambodia | 27 | 0.56 *(0.52–0.67)* | 0.22 | 209,779 *(195,794–251,735)* | 83,912 | 224,469 *(209,504–269,363)* | 89,788 |
| China | 70 | 0.30 *(0.23–0.37)* | 0.26 | 376,717 *(287,023–466,412)* | 322,900 | 244,030 *(185,928–302,133)* | 209,169 |
| Indonesia | 98 | 0.08 *(0.06–0.16)* | 0.30 | 302,430 *(226,822–604,860)* | 1,096,308 | 308,285 *(231,213–616,569)* | 1,117,532 |
| Laos | 16 | 0.62 *(0.44–0.81)* | 0.31 | 240,569 *(168,398–312,739)* | 120,284 | 171,966 *(120,376–223,556)* | 85,983 |
| Malaysia | 46 | 0.07 *(0.07–0.22)* | 0.22 | 61,873 *(61,873–206,244)* | 206,244 | 74,161 *(74,161–247,204)* | 247,204 |
| Myanmar | 43 | 0.33 *(0.30–0.51)* | 0.23 | 295,189 *(274,104–463,868)* | 210,849 | 214,624 *(199,293–337,266)* | 153,303 |
| Thailand | 235 | 0.06 *(0.03–0.11)* | 0.07 | 240,709 *(137,548–447,031)* | 292,290 | 233,216 *(133,266–433,115)* | 283,190 |
| Vietnam | 51 | 0.27 *(0.20–0.33)* | 0.10 | 241,146 *(172,247–292,820)* | 86,124 | 255,431 *(182,451–310,167)* | 91,225 |
| | **586** | | | **1,968,412** *(1,523,810–3,045,710)* | **2,418,912** | **1,726,182** *(1,336,193–2,739,372)* | **2,277,394** |

**Table S12 | Overlap between the area mapped as rubber here and protected areas[37] (PA) (all categories) and Key Biodiversity Areas[38] (KBA).**

| | Total mapped as rubber | Inside PA | Inside KBA | % in PA | % in KBA |
|---|---|---|---|---|---|
| Cambodia | 391,588 | 38,802 | 33,049 | 10% | 8% |
| China | 951,590 | 43,895 | 52,443 | 5% | 6% |
| Indonesia | 1,200,286 | 10,844 | 48,310 | 1% | 4% |
| Laos | 228,041 | 10,256 | 15,241 | 4% | 7% |
| Malaysia | 866,646 | 11,052 | 27,837 | 1% | 3% |
| Myanmar | 620,614 | 2,944 | 32,570 | 0% | 5% |
| Thailand | 3,435,939 | 140,643 | 140,299 | 4% | 4% |
| Vietnam | 848,900 | 76,158 | 32,711 | 9% | 4% |
| **Total** | **8,543,604** | **334,594** | **382,461** | **4%** | **4%** |

Draft - not peer reviewed; subject to revision. Findings are preliminary.

**Table 13 | Rubber area in Southeast Asia according to this and previous studies.** This is an extended version of Table 1 (with several rows repeating information from Table 1). Mapped areas are based on GAUL second-level administrative boundaries[39]. For China only the main production areas are included (Xishuangbanna and Hainan); mapped areas for Hainan exclude a small area where rubber was present.

Further sub-country mapping efforts not listed in the table reported 478 kha in 2020 in Xishuangbanna[40], 458 kha in 2021 in Hainan[41], and 592 kha (of monoculture rubber) in 2020 across Central, South and West Kalimantan[34].

**Sources for official rubber areas:** The areas were compiled from official sources reported in literature. Areas are not fully comparable as they refer to different plantation/age categories (e.g., mature, harvested, planted or unspecified rubber areas). Areas for Indonesia and Thailand refer to mature plantations in 2021, from BPS-Statistics Indonesia and the Thailand Office of Agricultural Economics, respectively, as reported by Hoang et al.[42]; for Vietnam to plantations harvested in 2021, from the General Statistics Office of Viet Nam, as reported by Hoang et al.[42]. For Malaysia the plantation type is unspecified; the area is for 2021 from the Malaysian Rubber Board and Department of Statistics Malaysia, as reported by Hoang et al.[42]. Areas for Laos and Myanmar refer to planted rubber area in 2018 and 2018/19, respectively, from the Lao Department of Forestry, as reported by Hoang et al.[42], and the Myanmar Department of Agriculture, as reported by Thu & Myint[43]. Areas for China are for 2022 and refer to planted area in Hainan from the Hainan Statistical Yearbook (2024), as reported by Liu et al.[41], and unspecified area in Xishuangbanna, from the National Bureau of Statistics of China (2023), as reported by Chen et al.[44]. The area for Cambodia refers to planted rubber area, including immature rubber, based on a newspaper report[45].

| **(kha)** | **Cambodia** | **China** | **Indonesia** | **Laos** | **Malaysia** | **Myanmar** | **Thailand** | **Vietnam** | **Total** |
|---|---|---|---|---|---|---|---|---|---|
| This study: error-adjusted estimate (mapped area) | 378<br>(392) | 1,256<br>(952) | 3,705<br>(1,200) | 385<br>(228) | 949<br>(867) | 907<br>(621) | 4,040<br>(3,436) | 878<br>(849) | **12,497**<br>(8,544) |
| Official rubber area | 404 | 813 | 3,776 | 275 | 1,137 | 659 | 3,915 | 931 | **11,915** |
| Wang et al.[26]: mapped area | 615 | 1,069 | 4,582 | 589 | 989 | 789 | 3,706 | 1,612 | **13,951** |
| Forest Data Partnership[28] model b: mapped area* | 751<br>985 | 1,392<br>1,583 | 90<br>113 | 2,163<br>3,049 | 491<br>563 | 3,315<br>4,675 | 5,687<br>6,615 | 3,303<br>4,330 | **17,192**<br>**21,914** |
| Forest Data Partnership[27] model a: mapped area | - | 957<br>1,258 | 2,692<br>4,837 | - | 396<br>703 | - | 3,048<br>5,099 | 2,281<br>3,732 | - |
| De Keersmaecker et al.[17]: mapped area | 413 | 534 | 1,301 | 868 | 263 | 1,148 | 2,952 | 657 | **8,135** |
| Chen et al.[46]: error adjusted estimate (mapped area) for 2022 | | | | | | | | 728<br>(754) | - |
| Xiao et al.[47]: estimate for 2024 | | | | 234 | | | | | - |

* The Forest Data Partnership (FDP) rubber probability models have been thresholded at values reported to balance precision and recall in two separate FDP evaluations: 0.59/0.44 and 0.23/0.35 for models 2025a/2025b, respectively. The thresholds result in over-mapping in Laos, Myanmar, and Vietnam, where according to our evaluations there is a risk of conflation with deciduous forests, and under-mapping in Indonesia, where detection power is low. As noted by the FDP authors, there is a need for local accuracy assessments to select the best thresholds.

| (kha) | Cambodia | China | Indonesia | Laos | Malaysia | Myanmar | Thailand | Vietnam | Total |
|---|---|---|---|---|---|---|---|---|---|
| Xiao et al.[30]: mapped area[†] for 2018 | 200 | 720[‡] | - | 700 | - | 680 | 4,650 | 740 | - |
| Petersen et al.[32]: mapped[§] for 2013-14 | 697 | - | 818 | - | 415 | - | - | - | - |
| Condro et al.[33]: mapped for 2019 | - | - | 299 | - | - | - | - | - | - |
| SDPT[48] V3: mapped for 2013-2019[**] | 858 | - | 1,115[††] | 686 | 421 | 603 | 4,223 | 587 | - |
| SDPT[29] V2: mapped for 2013-2019 | 866 | - | 1,115 | 706 | 421 | 603 | 4,250 | 591 | - |
| Hurni and Fox[49] estimated (mapped) for 2014[‡‡] | 2,974 (996) | 618 (323) | - | 765 (316) | - | 292 (125) | 2,861 (1,539) | 1,917 (984) | - |

[†] Here we use the areas reported by Xiao et al. (2021) in their manuscript's abstract.
[‡] Figure is for Yunan province.
[§] Petersen et al. also map mixed rubber areas; for comparability we only included pure rubber areas.
[**] Sources contributing to SDPT include Petersen et al. (2016) (for Malaysia, Indonesia, Cambodia), Xiao et al. (2021) (for Vietnam, Laos, Myanmar, Cambodia, Thailand), Condro et al. (2020) (for Indonesia) and Debonne et al. (2019) (for Cambodia). Both Petersen et al. (2016) and Debonne et al. (2019) also mapped rubber mixed with other crops; these areas were not included here for comparability. SDPT version 3 is under review and changes are still possible. In version 3, non-rubber datasets have been added for Cambodia, Laos, Myanmar, Thailand and Vietnam. These take precedence over Xiao et al. (2021) and Debonne et al. (2019), reducing rubber extent in areas of overlap (J. Richter, personal communication, 2026).
[††] Figure is approximate.
[‡‡] Areas for China only include Xishuangbanna, for Myanmar only Shan State, for Thailand only northeast Thailand, and for Vietnam only areas south of Hanoi.

**Table S14 | Types of remote sensing data used in the classifier.** For further details see Table S15, for variable importance in the classifier see Table S16, and for correlations between input variables Figure S9.

| Data source | Resolution | Time period | Usage in analysis |
|---|---|---|---|
| Sentinel-2 (Level 2A Surface Reflectance) | 10–20 m | 2021 (2019 for climatic zone B) | Annual band medians (visible, NIR, red-edge, SWIR); spectral indices (EVI, NDVI, NDWI, NBR, MNBR, SAVI); phenology metrics (defoliation/refoliation percentile based on NBR and NDVI, defoliation duration); NBR standard deviation at 20, 50, 100 m buffer sizes; NBR-based masks |
| Sentinel-1 GRD (C-band SAR) | 10 m | Annual time-series 2021 (2019 for climatic zone B) | VH cross-polarised band summary statistics (5th, 15th, 50th, 85th, 95th percentile) to capture canopy and phenological variation |
| PALSAR-2 annual mosaic (L-band SAR) | 25 m | Annual mosaic 2021 (2019 for climatic zone B) | HV cross-polarised band, providing longer-wavelength structural information to complement Sentinel-1 C-band |
| Canopy height[50] | 10 m | 2020 | Standard deviation of canopy height at 20, 50, and 100 m buffer sizes to distinguish even-aged plantations from natural forest |

Draft - not peer reviewed; subject to revision. Findings are preliminary.

**Table 15 | Variables used in the Random Forest classifier.** For details see Methods and Table S14.

| Variable | Category | Description |
|---|---|---|
| B2 | Optical bands | Sentinel-2 Level-2A surface reflectance blue band (490 nm) |
| B3 | | Sentinel-2 Level-2A surface reflectance green band (560 nm) |
| B4 | | Sentinel-2 Level-2A surface reflectance red band (665 nm) |
| B5 | | Sentinel-2 Level-2A surface reflectance red-edge 1 band (705 nm) |
| B6 | | Sentinel-2 Level-2A surface reflectance red-edge 2 band (740 nm) |
| B7 | | Sentinel-2 Level-2A surface reflectance red-edge 3 band (783 nm) |
| B8 | | Sentinel-2 Level-2A surface reflectance near-infrared (NIR) band (842 nm) |
| B8A | | Sentinel-2 Level-2A surface reflectance narrow near-infrared band (865 nm) |
| B11 | | Sentinel-2 Level-2A surface reflectance short-wave infrared 1 (SWIR1) band (1610 nm) |
| B12 | | Sentinel-2 Level-2A surface reflectance short-wave infrared 2 (SWIR2) band (2190 nm) |
| EVI | Spectral indices | Enhanced Vegetation Index derived from Sentinel-2 annual band medians |
| NDVI | | Normalized Difference Vegetation Index derived from Sentinel-2 annual band medians |
| NDWI_b8b11 | | Normalized Difference Water Index derived from Sentinel-2 bands B8 and B11 |
| SAVI | | Soil Adjusted Vegetation Index derived from Sentinel-2 annual band medians |
| NBR | | Normalized Burn Ratio derived from Sentinel-2 annual band medians |
| MNBR | | Modified Normalized Burn Ratio |
| NDVI_p15 | Phenology | 15th percentile of NDVI during defoliation window |
| NDVI_p85 | | 85th percentile of NDVI during refoliation window |
| NDVI_diff | | Difference between above NDVI percentiles to capture defoliation intensity |
| NBR_p15 | | 15th percentile of NBR during defoliation window |
| NBR_p85 | | 85th percentile of NBR during refoliation window |
| NBR_diff | | Difference between above NBR percentiles to capture defoliation intensity |
| defol_day | | Day of defoliation onset |
| defol_dur | | Duration of the defoliation period |
| refol_day | | Day of refoliation onset |
| NBR_tex20 | Texture | Standard deviation of NBR within a 20 m buffer window |
| NBR_tex50 | | Standard deviation of NBR within a 50 m buffer window |
| NBR_tex100 | | Standard deviation of NBR within a 100 m buffer window |
| canopyHeight_20 | | Standard deviation of canopy height within a 20 m buffer window |
| canopyHeight_50 | | Standard deviation of canopy height within a 50 m buffer window |
| canopyHeight_100 | | Standard deviation of canopy height within a 100 m buffer window |
| VH | Radar | Median of Sentinel-1 VH cross-polarised band |
| VH_p5 | | 5th percentile of Sentinel-1 VH band |
| VH_p15 | | 15th percentile of Sentinel-1 VH band |
| VH_p85 | | 85th percentile of Sentinel-1 VH band |
| VH_p95 | | 95th percentile of Sentinel-1 VH band |
| VH_stdDev | | Standard deviation of Sentinel-1 VH band |
| HV | | Median of PALSAR-2 HV cross-polarised band |
| latitude | Other | Geographic latitude coordinate |
| longitude | | Geographic longitude coordinate |

**Table S16 | Variable importance in the Random Forest classifier for climatic zone A and B.** For row key see Table S15 and for correlations between these input variables Figure S9.

| | A | B |
|---|---|---|
| **B11** | 76.18 | 51.06 |
| **B12** | 70.87 | 35.34 |
| **B2** | 41.6 | 28.58 |
| **B3** | 41.78 | 28.4 |
| **B4** | 49.52 | 29.03 |
| **B5** | 55.44 | 29.84 |
| **B6** | 66.36 | 32.23 |
| **B7** | 52.96 | 28 |
| **B8** | 61.03 | 26.54 |
| **B8A** | 60.82 | 18.22 |
| **EVI** | 52.6 | 41.33 |
| **HV** | 57.85 | 68.93 |
| **MNBR** | 56.84 | 30.96 |
| **NBR** | 50.82 | 35.37 |
| **NBR_diff** | 124.62 | 32.09 |
| **NBR_p15** | 93.77 | 32.9 |
| **NBR_p85** | 55.75 | 30.99 |
| **NBR_tex100** | | 30.6 |
| **NBR_tex20** | 62.65 | 48.61 |
| **NBR_tex50** | 60.75 | 33.28 |
| **NDVI** | | 27.37 |
| **NDVI_diff** | 61.46 | 31.72 |
| **NDVI_p15** | 51.07 | 29.23 |
| **NDVI_p85** | 57.14 | 36.61 |
| **NDWI_b8b11** | 47.55 | 34.79 |
| **SAVI** | 50.5 | 28.37 |
| **VH** | 48.77 | 31.22 |

| | A | B |
|---|---|---|
| **VH_p15** | 52.05 | 26.32 |
| **VH_p5** | 43.47 | 30.13 |
| **VH_p85** | 38.02 | 27.42 |
| **VH_p95** | 31.61 | 27.66 |
| **VH_stdDev** | 44.41 | 29.12 |
| **canopyHeight_100** | 43.51 | 25.87 |
| **canopyHeight_20** | 31.84 | 22.3 |
| **canopyHeight_50** | 41.21 | 29.03 |
| **defol_day** | 75.13 | 29.81 |
| **defol_dur** | 64.15 | 27.6 |
| **latitude** | 138.54 | 40.05 |
| **longitude** | 94.87 | 40.87 |
| **refol_day** | 80.28 | 41.32 |

**Table S17 | Interpretation criteria for classifying land cover prior to rubber.** Over 90% of points were assessed by at least two interpreters (and over 95% of points in cases of potential clearances). Points with conflicting interpretation were reviewed again by both interpreters, in difficult cases with the help of a third interpreter. Where available, interpreters used very high-resolution imagery in Google Earth Pro, in addition to lower-resolution (Sentinel 2, Landsat) time series imagery and auxiliary information such as local reports of land cover histories, and other spatial layers (for example, for protected areas[37], Key Biodiversity areas[38] and primary humid forest[14]).

Agroforests, community woodlots and other intermediate forms of tree cover were the most difficult to classify. Depending on their degree of naturalness and the characteristics of the plantation replacing them, their conversion may either be largely inconsequential or constitute a loss of semi-natural tree cover. We attempted to reflect this context in our interpretation. For example, we assigned points to 'semi-natural' when (seemingly) mostly semi-natural vegetation was converted to large industrial monocultures, but we assigned points to already 'modified' when there was no obvious or substantial change in the degree of naturalness and/or structural diversity (e.g. small plantations replacing small agroforests). However, in practice, many sample points with such difficult-to-classify tree covers had to be classified as "unknown" as image quality was often insufficient.

| Prior landcover | Description | Certainty | Criteria | Potential errors |
|---|---|---|---|---|
| **Natural tree cover** | Naturally regenerating tree cover with little evidence for human management/influence | 1 | At least one high-resolution image shows natural tree cover immediately prior to rubber cultivation, and all other evidence is consistent with that interpretation (for example, there are no signs of roads or settlements, there are local reports of deforestation, the clearance is often visible on high-resolution imagery, e.g. with an image showing fallen stems, and/or the area is inside a protected area or was classed as primary humid forest in 2001). | Overgrown and /or long-rotation plantations, secondary forests, and agroforests, may have been misclassified as natural tree cover. |
| | | 0.95 | At least one high-resolution image shows natural tree cover prior to rubber cultivation, and all other evidence is consistent with that interpretation, but with slightly less certainty than above. | |
| | | 0.9 | At least one high-resolution image shows natural tree cover prior to rubber cultivation but with less certainty, and/or there is no high-resolution image of the area itself but an adjacent high-resolution image shows prior natural tree cover, and low-resolution imagery suggests that this tree cover extended into the target area. | |
| | | 0.85 | At least one high-resolution image suggests that there was natural tree cover, but with low certainty (for example, the area is smaller and could also be semi-natural). | |
| | | 0.8 | High-resolution images appear to show natural tree cover, but the evidence is less conclusive than above. For example, the area is smaller, there are nearby settlements and/or fields - suggesting this could also be semi-natural or even planted tree cover. Alternatively, only lower-resolution | |

| | | | | |
|---|---|---|---|---|
| | | | imagery is available, but several pieces of evidence point into the direction of natural tree cover (e.g. no obvious prior tree rotation and/or the area was classed as primary humid forest in 2001). | |
| | | 0.75 | Very difficult to determine. At least one high-resolution image appears to show natural tree cover but this cannot be identified with certainty. Alternatively, only lower-resolution imagery is available – typically showing a large, dark green patch (with no visible roads, settlements, or plantation structure), followed by a sudden, large-scale clearance (shift from green to yellow/brown). | |
| **Semi-natural tree cover** | Predominantly naturally regenerating tree cover influenced by past and/or present human activity, including secondary, logged, grazed or fragmented forests, as well as smaller forest patches embedded within mosaics of bush and agricultural land, or located near roads and settlements | *As above* | As for natural tree cover but there is evidence of prior clearance, human use and/or degradation of the tree cover replaced by rubber. For example, areas may be fragmented, criss-crossed by paths, located near roads or settlements, or embedded within mosaics of fields and secondary forest. Prior disturbance may also be indicated by yellow-brown colours in low-resolution imagery or by sparse or bush-like tree cover in high-resolution imagery in areas where closed-canopy forest is the potential natural vegetation.<br><br>Add a flag to areas with signs of heavy degradation. These include areas where scrub may have replaced original tree cover or heavily fragmented and/or small forest patches of only a few hectares in size. | The degree of naturalness in partially modified landscapes was particularly difficult to determine. Overgrown plantations may have been misclassified as semi-natural tree cover, whereas naturally sparse tree cover, bushland or savannahs may have been misinterpreted as degraded semi-natural tree cover. |
| **Modified** | Areas already modified prior to 1990, including non-forested areas and/or plantations | 1 | High-resolution images show no tree cover or a plantation prior to rubber cultivation, and/or local reports confirm the prior existence of a plantation. If only low-resolution imagery is available, it shows structural features typical of plantations (terracing, grid patterns, or dense road networks) or colours indicative of prior modification (for example typical earth/field tones or bright green, often seen in rubber plantations). | Small-scale clearances of (semi-)natural tree cover may have been missed, particularly where only low-resolution imagery was available. Furthermore, losses of secondary or deciduous forest may have been missed, as they can resemble cultivated landscapes (particularly on lower resolution imagery). Landscape context was not always a reliable indicator as deciduous and secondary forests often occur within areas of agriculture. |
| | | 0.95 | Similar to above, but the evidence is less clear - for example, there is no high-resolution image, or plantation or clearance features on low-resolution imagery are less distinct. | |
| | | 0.9 | Low-resolution images show diverse colours (yellow, brown, green) or frequent colour changes over time, suggesting a modified (cultivated) landscape | |

Draft - not peer reviewed; subject to revision. Findings are preliminary.

| | | | |
|---|---|---|---|
| | | 0.8 | Similar to above but evidence is less clear |
| | | 0.75 | Very difficult to determine. The area itself may appear forested (dark green), but surrounding features such as roads and plantations suggest that this could be a plantation. Alternatively, there is no visible clearance before rubber planting and/or high-resolution imagery shows swidden agriculture with the area itself appearing more like a plantation than like a forest. |
| **Unknown** | | | Prior landcover cannot be determined – for example because there is no high-resolution imagery and lower-resolution imagery is inconclusive. Alternatively, higher-resolution images are available, but it is not clear whether the images show natural, planted or semi-natural tree cover.<br><br>(This category was also assigned when interpreters could not reach consensus.) |

### Supporting figures

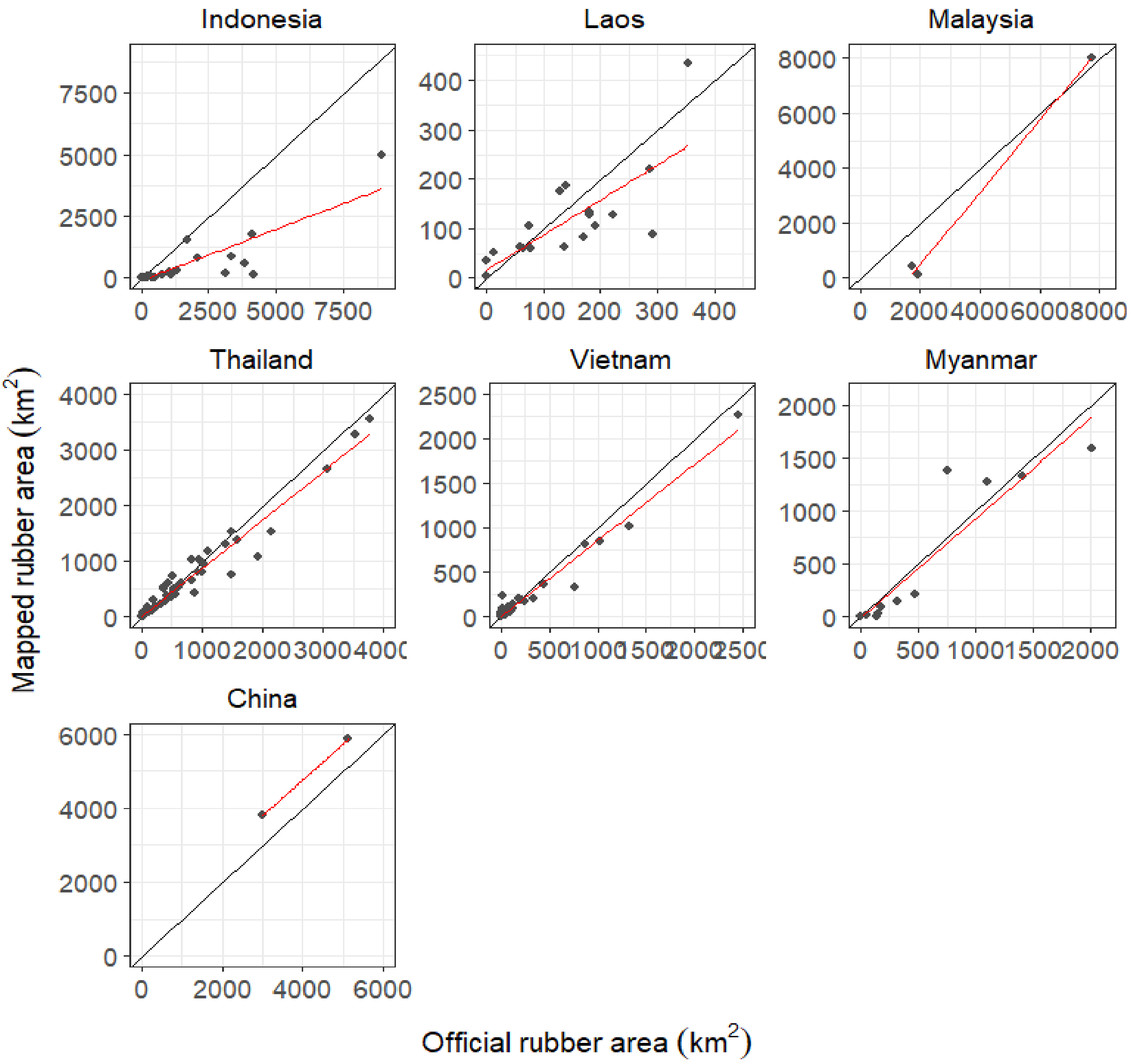

**Figure S1 | Relationship between official data and areas mapped as rubber in this study at subnational level.** While there was broad agreement between officially reported rubber areas and rubber areas mapped in this study, our map tends to under-map rubber extent (particularly in Indonesia). Exceptions are China, where we mapped more rubber than officially reported, and Myanmar and Laos, where, in addition to mapping more rubber, our mapped areas did not always align with officially reported areas.

Sources of official data are reported in Table S13.

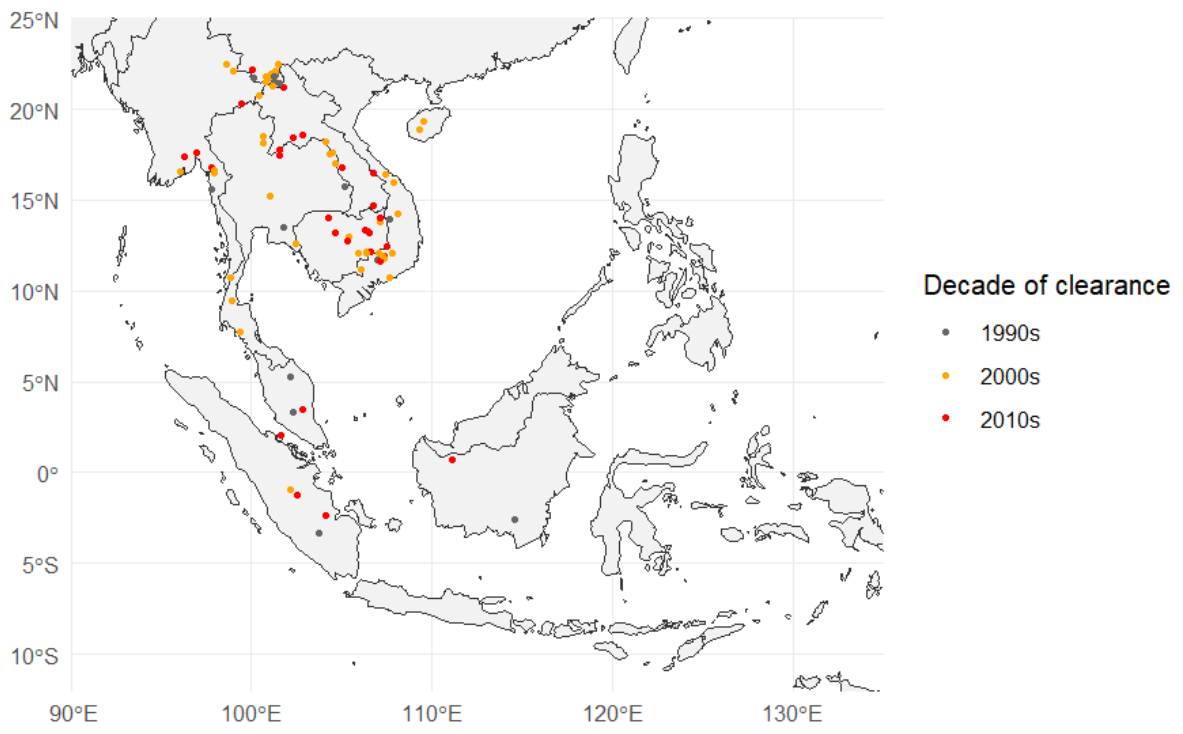

**Figure S2 | Sample points for which the pre-rubber landcover was natural or semi-natural tree cover (by decade of clearance).** There is a likely bias towards more recent clearances, as, with satellite imagery improving, post 2010-clearances were a lot easier to detect than clearances in the 1990s (Fig. S3). This bias will be particularly pronounced in equatorial regions due to the added challenge of cloud and haze there.

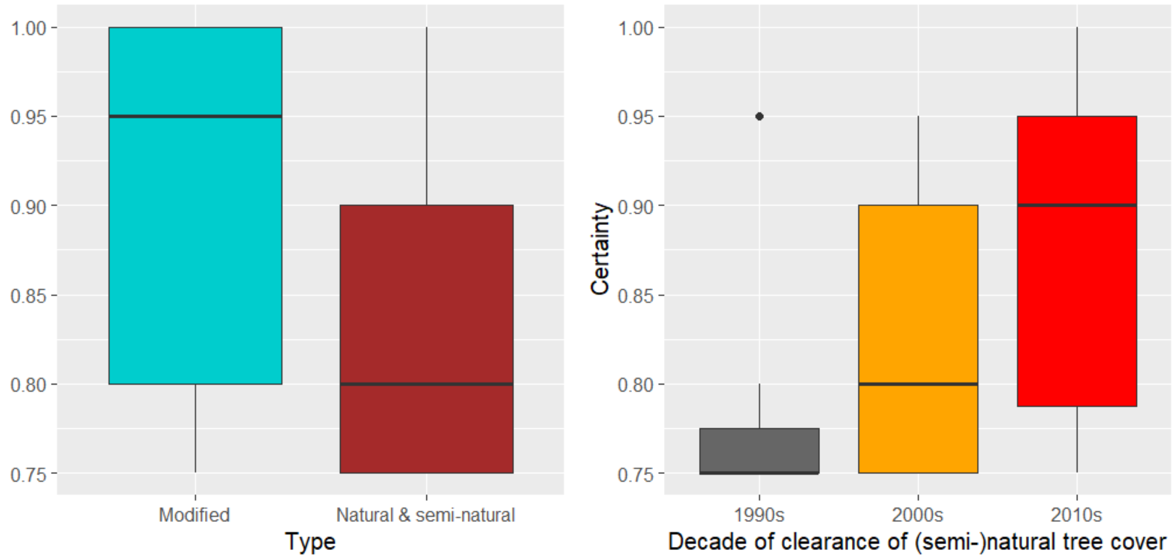

**Figure S3 | Certainty of pre-rubber land cover classifications (based on subjective scores assigned by interpreters).** Left: Certainty with which interpreters felt able to identify the pre-rubber land cover type. Right: Certainty with which interpreters felt able to identify clearances across decades. For details on how interpreters assigned certainty scores see Table S17.

Modified landscapes could be detected with higher certainty than (semi-)natural ones. This is because modified landscapes can have distinctive features visible even on low-resolution imagery (grid or terraced planting patterns, dense road networks, and yellow or brown colours characteristic of fields and clearings). Identifying natural tree cover on low-resolution images was generally only possible if it formed part of large contiguous evergreen forests. Smaller patches of natural forests, particularly deciduous, disturbance-adapted and/or secondary, were difficult to identify on low-resolution imagery as they can resemble agricultural rotation. There may thus be a bias towards underestimating losses these forest types and/or for time periods and areas lacking clear high-resolution images (often prior to 2000 and in countries near the equator).

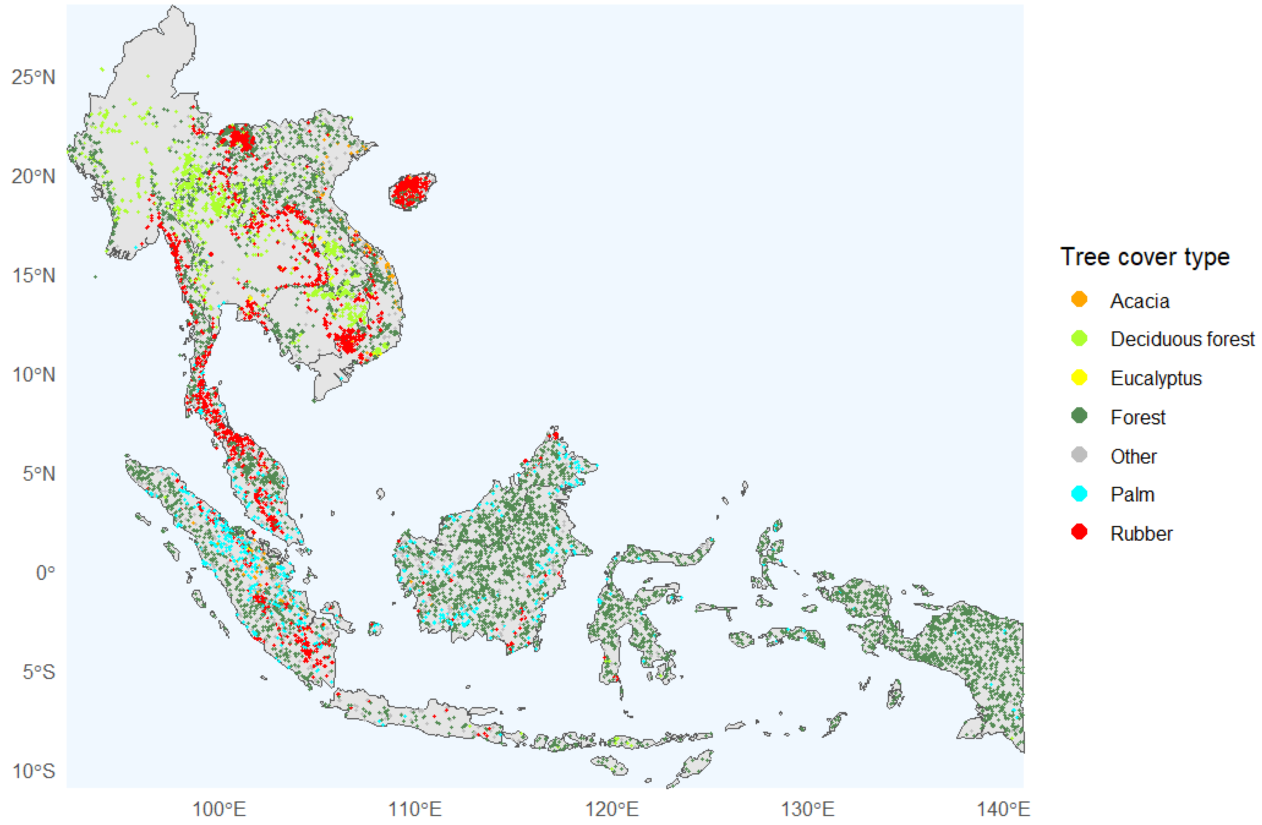

**Figure S4 | Training sample points.** Training points (in total 12,499) covered rubber (3,578 points); forest (7,094 points for deciduous, evergreen, degraded, and primary forest types); palm plantations (608 points); *Acacia* (170 points) and *Eucalyptus* (80 points) plantations; other tree plantations and land covers (919 points, including for cashew, mango, coffee, teak, shrubs, riverine vegetation, mangroves, and floodplains).

The rubber classification algorithm was trained on 10,143 points (3,345 for rubber and 6,798 for other tree cover). An additional 2,036 points were collected for analyses presented in Tables S7-S8.

Data sources: We used 3,826 data points from Wang et al.[26], who compiled reference data from multiple sources, including data contributed by the World Agroforestry Centre, Yang et al.[51], Lan et al.[52] and Hurni & Fox[49]. Over 50% of these points had originally been based on field data. The remaining 8,623 points were collected specifically for this study from very high-resolution imagery in Google Earth Pro, with the interpretation supported, where available, by Google Street View, polygons and other data shared by rubber supply chain actors, and information available on the internet about industrial rubber estates and other plantations. In more uncertain cases we also used time-series imagery from Sentinel-2 and Landsat 5, 7, and 8. Because a substantial proportion of the data was derived through visual interpretation of imagery, misclassifications may have occurred; for example, some agroforestry systems may have been classified as forest given structural and spectral similarity in remotely sensed images.

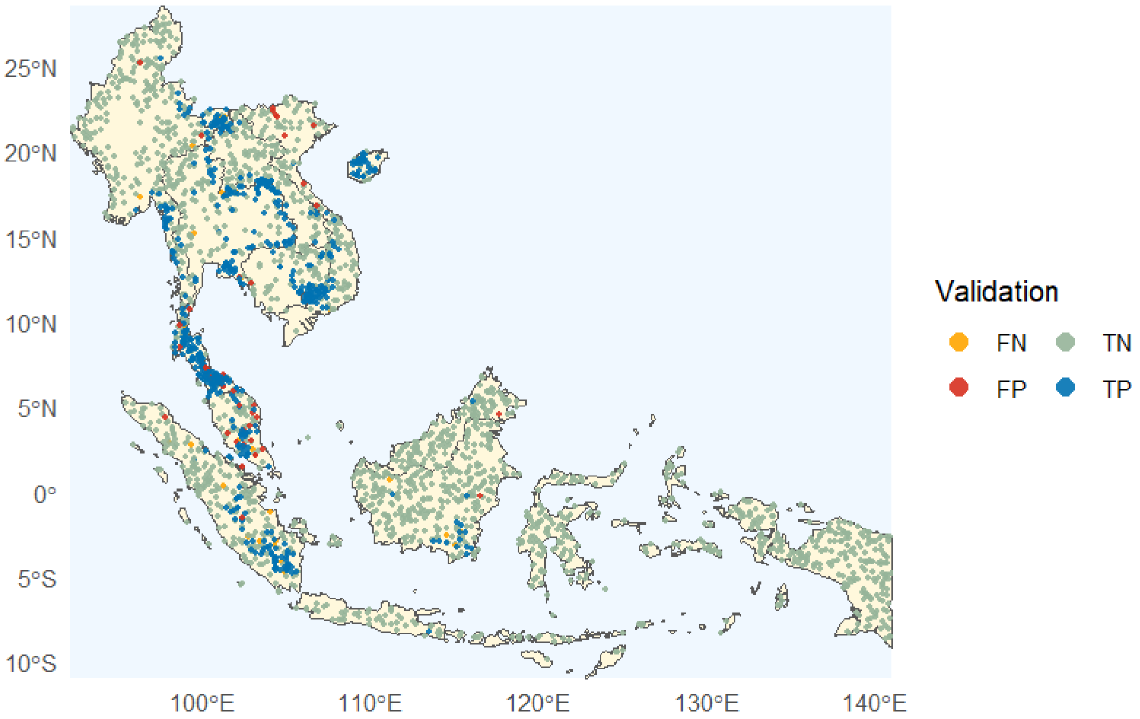

**Figure S5 | Validation points.** The 2,500 points were generated using the *stratifiedSample* function in Google Earth Engine[53] (600 points for areas mapped as rubber and 1,900 for areas mapped as other tree cover). As above in Fig. S4, where available, the interpretation was supported by auxiliary information from Google Street View, data contributed by stakeholders, information available on the internet, and satellite image time series. The validation data points were generally assessed by at least two, and in cases of uncertain interpretation, three interpreters.

**Legend key:** TP = true rubber positives (*n* = 556), TN = true rubber negatives (*n* = 1,870), FP = false rubber positives (*n* = 44), FN = false rubber negatives (*n* = 30).

For full validation results see Tables S1, S3 and S4, and for a description of false positives Table S5. With the classification mostly based on visual interpretation of imagery, misclassifications may have occurred. For instance, agroforestry systems may have been classified as non-rubber because of their structural and spectral similarity to natural forest in remotely sensed images.

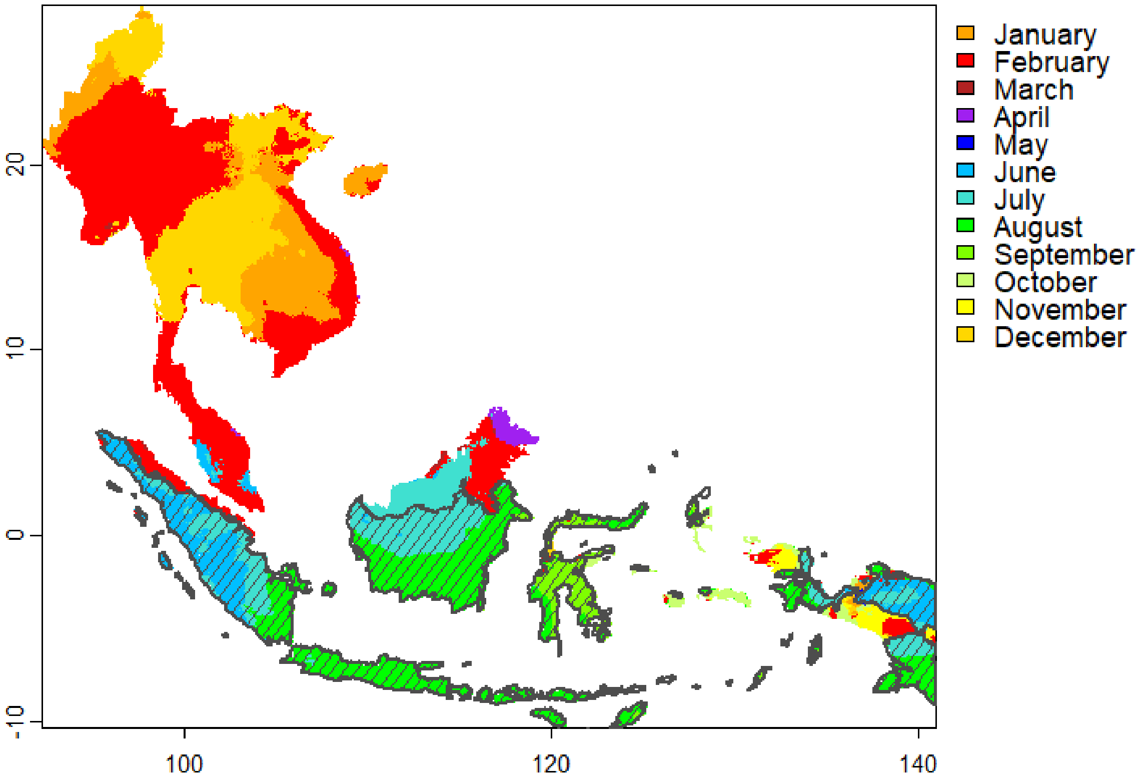

**Figure S6 | Month of minimum precipitation across Southeast Asia.** Colours indicate the month in which mean monthly precipitation was lowest according to a multisource dataset[54] based primarily on SM2RAIN-ASCAT rainfall data (2007–2018). The outlined and hatched region shows climatic zone B, where rubber defoliation was assumed to occur between June and September. In the rest of the study area, defoliation was assumed to occur between January and February.

As shown here and in Fig. S8, the broad climatic zones do not fully capture the complex dry-season patterns in Malaysia and Indonesia. While this could be refined using field observations of rubber phenology, consistently high moisture levels (Fig. S7), together with cloud and haze, often weaken the optical defoliation signal in these regions. Refinements of climatic zones may therefore not significantly improve rubber detection. This is also indicated by the finding that in zone B the classification algorithm relied more strongly on radar and textural metrics than phenology information (Table S16).

Instead, a step-change in detection is more likely to come from more field-based rubber data and algorithms being run locally, as exemplified by the significantly higher rubber detection of the map by Le Maire et al.[34].

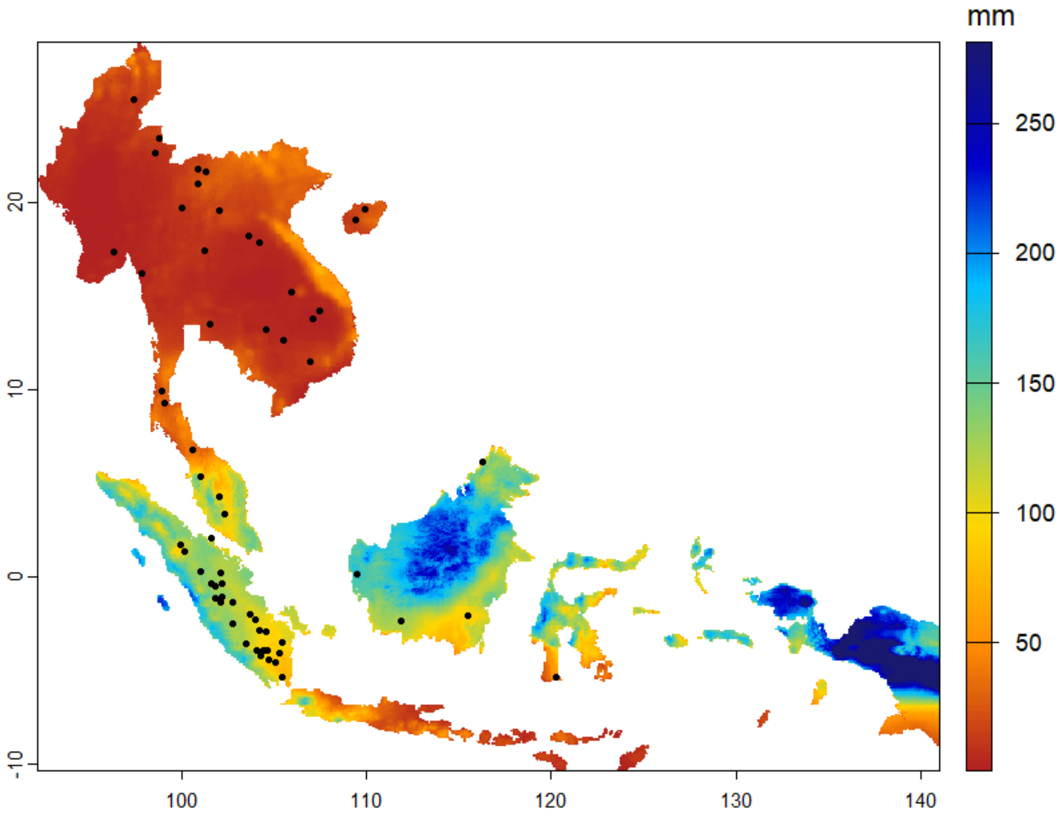

**Figure S7 | Spatial distribution of minimum monthly precipitation across Southeast Asia.** Values indicate precipitation totals during the driest month of the year, derived from a multisource dataset[54] primarily based on SM2RAIN-ASCAT rainfall data (2007–2018). Most areas in Malaysia and Indonesia have a more humid and less seasonal climate with comparatively high precipitation levels even during the driest month(s).

The black dots are random rubber samples used to generate Fig. S8.

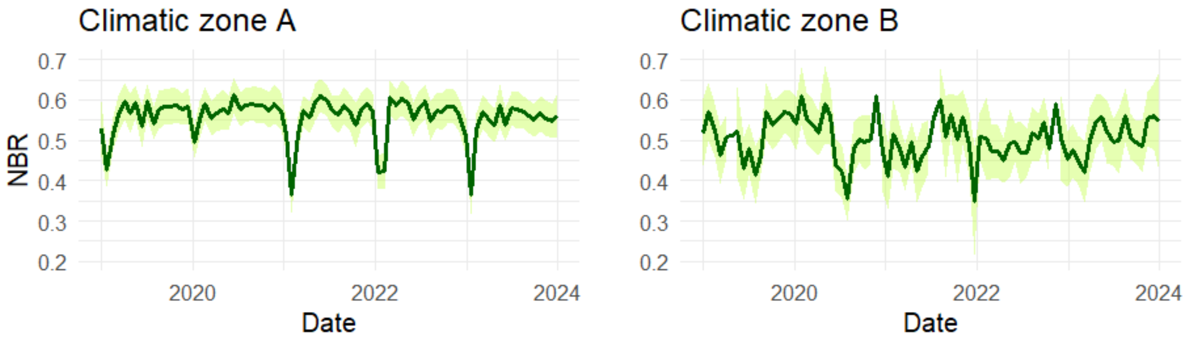

**Figure S8 | Seasonal dynamics in rubber canopy phenology in climatic zones A and B.** The time series shows Normalized Burn Ratio (NBR) values for 30 randomly selected rubber points (Fig. S7) within each climatic zone. The solid line is a locally weighted regression (LOESS) fit to the mean of observations (span = 0.01; 95% confidence interval shaded in lighter green).

In climatic zone A, the NBR time series showed a pronounced seasonal drop associated with dry-season rubber defoliation in January-February. However, the target mapping year (2020) was characterised by wetter than average conditions, reducing the strength of the defoliation signal. We therefore used imagery from 2021 to map rubber in zone A.

In zone B the drop in NBR was more variable and difficult to predict. This reflects both ground conditions and limitations of optical imagery: when soil moisture does not drop below critical levels, leaf renewal in rubber trees can may be more gradual, less synchronous and/or entirely absent. In addition, image gaps and remnant cloud, haze and cloud shadow not detected by cloud and shadow masks render the time series noisy and sparse. After examining NBR time series for rubber and various other tree covers it can be conflated with, and numbers of available images, we identified 2019 as the best mapping year for zone B.

Draft - not peer reviewed; subject to revision. Findings are preliminary.

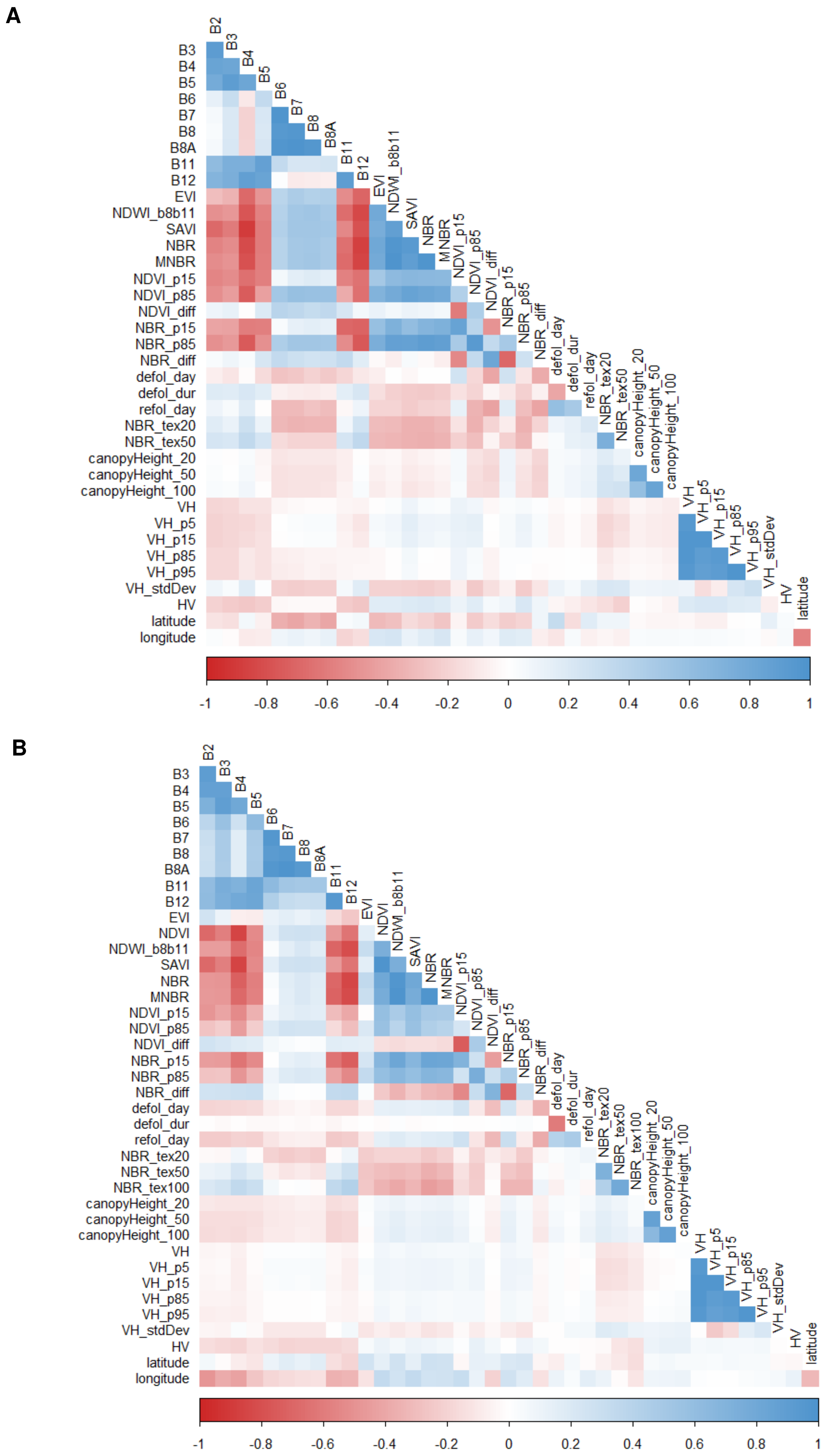

**Figure S9 | Correlations (Spearman) between classifier input variables in climatic zones A (top panel) and B (bottom panel).**

Draft - not peer reviewed; subject to revision. Findings are preliminary.

Draft - not peer reviewed; subject to revision. Findings are preliminary.

24 Vancutsem, C. *et al.* Long-term (1990–2019) monitoring of forest cover changes in the humid tropics. *Science Advances* **7**, eabe1603 (2021). https://doi.org/doi:10.1126/sciadv.abe1603
25 Sheil, D., Descals, A., Meijaard, E. & Gaveau, D. Rubber planting and deforestation. *Nature* **644**, E13-E19 (2025). https://doi.org/10.1038/s41586-025-08847-w
26 Wang, Y. *et al.* High-resolution maps show that rubber causes substantial deforestation. *Nature* **623**, 340-346 (2023). https://doi.org/10.1038/s41586-023-06642-z
27 Forest Data Partnership. Rubber probability model 2025a. (2025).
28 Forest Data Partnership. Rubber probability model 2025b. (2025).
29 Richter, J. *et al.* Spatial Database of Planted Trees (SDPT Version 2.0). (World Resources Institute, Washington, DC, 2024).
30 Xiao, C. *et al.* Latest 30-m map of mature rubber plantations in Mainland Southeast Asia and Yunnan province of China: Spatial patterns and geographical characteristics. *Progress in Physical Geography: Earth and Environment* (2021). https://doi.org/10.1177/0309133320983746
31 Debonne, N., van Vliet, J. & Verburg, P. Future governance options for large-scale land acquisition in Cambodia: Impacts on tree cover and tiger landscapes. *Environmental Science & Policy* **94**, 9-19 (2019). https://doi.org/10.1016/j.envsci.2018.12.031
32 Petersen, R. *et al.* Mapping Tree Plantations with Multispectral Imagery: Preliminary Results for Seven Tropical Countries. (World Resources Institute, Washington, DC, 2016).
33 Condro, A., Setiawan, Y., Prasetyo, L., Pramulya, R. & Siahaan, L. Retrieving the National Main Commodity Maps in Indonesia Based on High-Resolution Remotely Sensed Data Using Cloud Computing Platform. *Land* **9** (2020). https://doi.org/10.3390/land9100377
34 le Maire, G. *et al.* Mapping rubber monoculture and jungle rubber across Kalimantan using multisource satellite data. *International Journal of Remote Sensing*, 1-25 (2025). https://doi.org/10.1080/01431161.2025.2545024
35 Wohlfart, C., Wegmann, M. & Leimgruber, P. Mapping Threatened Dry Deciduous Dipterocarp Forest in South-East Asia for Conservation Management. *Tropical Conservation Science* **7**, 597-613 (2014). https://doi.org/10.1177/194008291400700402
36 Kennedy, R. *et al.* Implementation of the LandTrendr Algorithm on Google Earth Engine. *Remote Sensing* **10** (2018). https://doi.org/10.3390/rs10050691
37 UNEP-WCMC and IUCN. The World Database on Protected Areas (WDPA). (Cambridge. Available at: www.protectedplanet.net, 2025).
38 BirdLife International. World Database of Key Biodiversity Areas. Developed by the KBA Partnership: BirdLife International, International Union for the Conservation of Nature, American Bird Conservancy, Amphibian Survival Alliance, Conservation International, Critical Ecosystem Partnership Fund, Global Environment Facility, Re:wild, NatureServe, Rainforest Trust, Royal Society for the Protection of Birds, Wildlife Conservation Society and World Wildlife Fund. March 2022 version. Available at http://keybiodiversityareas.org/kba-data/request. (2022).
39 Food and Agriculture Organization of the United Nations (FAO). (Available at Google Earth Engine, 2015).
40 Li, Y., Liu, C., Zhang, J., Zhang, P. & Xue, Y. Monitoring Spatial and Temporal Patterns of Rubber Plantation Dynamics Using Time-Series Landsat Images and Google Earth Engine. *IEEE Journal of Selected Topics in Applied Earth Observations and Remote Sensing* **14**, 9450-9461 (2021). https://doi.org/10.1109/jstars.2021.3110763
41 Liu, X., Liao, J., Jing, R., Ye, H. & Teng, L. Extraction of Rubber Plantations on Hainan Island, China, Using Multi-Source Remote Sensing Images During 2021–2025. *Forests* **16** (2025). https://doi.org/10.3390/f16121773
42 Hoang, N. T., Potapov, P., Olofsson, P. & Kanemoto, K. Accuracy of rubber-related deforestation maps. *Nature* **644**, E13-E19 (2025). https://doi.org/10.1038/s41586-025-08847-w
43 Thu, K. M. & Myint, T. Myanmar's Natural Rubber Forward Policy: How to do it? , (2020).
44 Chen, Y. *et al.* Spatiotemporal trends of rubber defoliation and refoliation and their responses to abiotic factors in the northern edge of the Asian tropics. *Industrial Crops and Products* **226** (2025). https://doi.org/10.1016/j.indcrop.2025.120753
45 Khmer Times. in *Khmer Times* (2022).
46 Chen, B. *et al.* A full time series imagery and full cycle monitoring (FTSI-FCM) algorithm for tracking rubber plantation dynamics in the Vietnam from 1986 to 2022. *ISPRS Journal of Photogrammetry and Remote Sensing* **220**, 377-394 (2025). https://doi.org/10.1016/j.isprsjprs.2024.12.018

47 Xiao, C. *et al.* RPI-GMM: A novel structure-based and phenology-independent algorithm for mapping latest 10-m resolution national-level rubber plantations. *Remote Sensing of Environment* **334**, 115241 (2026). https://doi.org/https://doi.org/10.1016/j.rse.2026.115241

48 Richter et al. Spatial Database of Planted Trees (SDPT Version 3.0). (World Resources Institute, Washington, DC, in review).

49 Hurni, K. & Fox, J. The expansion of tree-based boom crops in mainland Southeast Asia: 2001 to 2014. *Journal of Land Use Science* **13**, 198-219 (2018). https://doi.org/10.1080/1747423x.2018.1499830

50 Lang, N., Jetz, W., Schindler, K. & Wegner, J. D. A high-resolution canopy height model of the Earth. *Nature Ecology & Evolution* **7**, 1778-1789 (2023). https://doi.org/10.1038/s41559-023-02206-6

51 Yang, J., Xu, J. & Zhai, D.-L. Integrating Phenological and Geographical Information with Artificial Intelligence Algorithm to Map Rubber Plantations in Xishuangbanna. *Remote Sensing* **13** (2021). https://doi.org/10.3390/rs13142793

52 Lan, G. *et al.* Main drivers of plant diversity patterns of rubber plantations in the Greater Mekong Subregion. *Biogeosciences* **19**, 1995-2005 (2022). https://doi.org/10.5194/bg-19-1995-2022

53 Gorelick, N. *et al.* Google Earth Engine: Planetary-scale geospatial analysis for everyone. *Remote Sensing of Environment* **202**, 18-27 (2017). https://doi.org/10.1016/j.rse.2017.06.031

54 Hengl, T. & Parente, L. (Zenodo: https://doi.org/10.5281/zenodo.6458580, 2022).
